# Spatial and single-cell transcriptomics reveal neuron-astrocyte interplay in long-term memory

**DOI:** 10.1101/2023.03.20.533566

**Authors:** Wenfei Sun, Zhihui Liu, Xian Jiang, Michelle B. Chen, Hua Dong, Jonathan Liu, Thomas C. Südhof, Stephen R. Quake

## Abstract

Memory encodes past experiences, thereby enabling future plans. The basolateral amygdala (BLA) is a center of salience networks that underlie emotional experience and thus plays a key role in long-term fear memory formation^1, 2^. Here we used spatial and single-cell transcriptomics to illuminate the cellular and molecular architecture of the role of the basolateral amygdala in long-term memory. We identified transcriptional signatures in subpopulations of neurons and astrocytes that were memory-specific and persisted for weeks. These transcriptional signatures implicate neuropeptide signaling, mitogen-activated protein kinase (MAPK), brain-derived neurotrophic factor (BDNF), cAMP response element-binding protein (CREB), ubiquitination pathways, and synaptic connectivity in long-term memory. We also discovered that a neuronal sub-population, defined by increased *Penk* expression and decreased *Tac* expression, constitutes the most prominent component of the BLA’s memory engram. These transcriptional changes were observed both with single-cell RNAseq and with single-molecule spatial transcriptomics in intact slices, thereby providing a rich spatial map of a memory engram. The spatial data enabled us to show that this neuronal subpopulation further interacts with spatially related astrocytes that are essential for memory consolidation, indicating that neurons require interactions with astrocytes to encode long term memory.

## Introduction

Consolidation of newly acquired memories into long-term memories and reconsolidation of memory during recall require transcription and translation, as shown by extensive studies of the role of gene expression during learning and memory^3–5^. Although key transcription factors in learning and short term memory, such as CREB^6–9^, have been identified, the overall nature of long-term memories, which can persist a lifetime, remains enigmatic.^10^ Gene expression changes are known to be essential for long-term memory, but the cell types and the nature of the transcriptional programs involved are incompletely understood. Moreover, multiple brain regions have been implicated in long-term memory formation and storage but it is unknown whether similar transcriptional processes are used in different regions of the brain.

Here, we performed high-resolution spatial and single-cell transcriptomics to comprehensively analyze the changes in the transcriptomic landscape during long-term memory formation in mice. We identified memory-specific gene expression changes in the amygdala, which is a complex brain region whose basolateral area (BLA) is implicated in short- and long-term memories associated with salient experiences, such as fear. Lesions of the BLA abolish both short- and long-term fear memory.^11^ In fear learning paradigms, suppressing RNA transcription in the BLA before training attenuates fear memory consolidation without impacting the freezing response to a foot shock.^12^ Inhibiting protein synthesis in the BLA immediately after training^13^ or after reactivation^14^ also impairs long-term memory consolidation, but does not impact short-term memory recall.^14^

Our results show that neurons and astrocytes in the BLA exhibit memory-specific persistent transcriptional signatures that correspond to multiple signaling pathways but are highly specific to a small subset of cells representing engram cells. We identified a sub population of neuronal cells with increased *Penk* expression and decreased *Tac* expression (P+T- neurons) which are the most prominent part of the long-term memory engram. Using the spatial transcriptomic data, we discovered that there is a population of astrocytes which are co-located with the P+T- neurons, which undergo gene expression changes in forming long term memory, and which are required for long term memory consolidation. Finally, integration of this data with previous data^15^ on long-term contextual fear memory in the medial prefrontal cortex enabled us to examine region-specific versus general gene expression changes. These results show that similar molecular programs and cell types are used in long-term fear memories across both regions of the brain.

## Spatial and deep single-cell transcriptomics probing persistent transcriptional changes during fear-memory recall

In TRAP2 mice, cellular activation induces expression of tamoxifen-dependent Cre-ERT2 recombinase embedded in the *Fos* gene. As a result, TRAP2 mice crossed to Ai14 tdTomato (tdT) reporter mice express tdT only if they are both stimulated and exposed to tamoxifen^16^. We trained TRAP2 mice crossed to Ai14 mice by fear conditioning on day 0 and triggered recall of long-term fear memories by returning the mice to the training context 16 days later with simultaneous injection of tamoxifen. We then analyzed the amygdala of the mice on day 25, nine days after recall, by spatial transcriptomics and full-length deep single-cell RNAseq (Fig. 1**a**,**b**). As controls for this <u>F</u>ear training and <u>R</u>ecall (FR) condition, we used <u>H</u>ome <u>C</u>age (‘HC’) mice that were left in their home cage, and <u>N</u>o <u>F</u>ear (‘NF’) and <u>N</u>o <u>R</u>ecall (‘NR’) mice that were exposed to all manipulations except that they received either no electrical shock during training (NF) or were not exposed to the recall condition (NR). The goal of this experimental design was to mark engram cells that are activated during the recall and become tdT+, allowing us to identify fear-specific memory genes that are differentially expressed in these engram cells and are not induced by salience only (the NF condition).

**Fig. 1.**
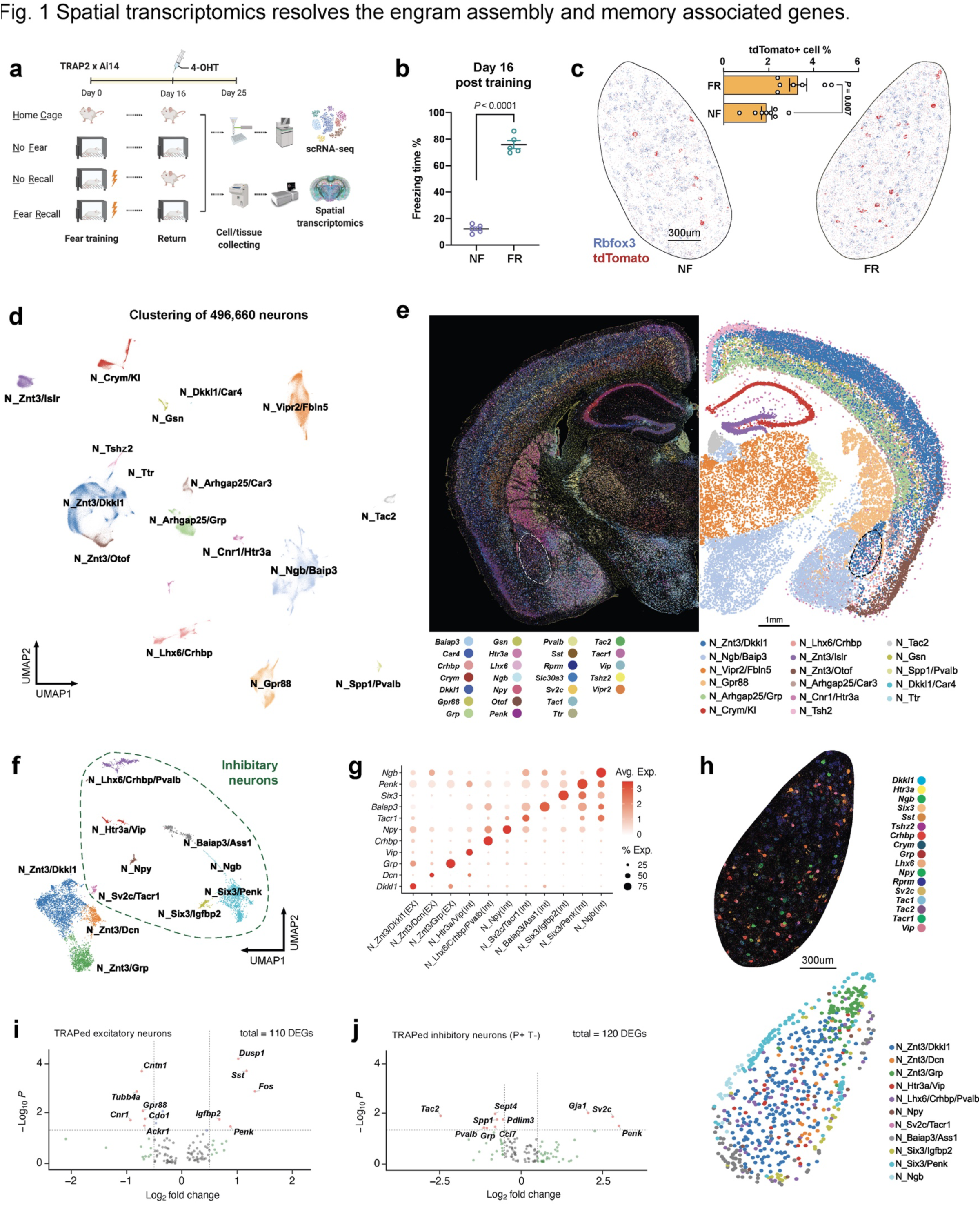
Spatial transcriptomics resolves the engram assembly and memory associated genes. **a)** Experiment scheme for tracing engram cells in a fear conditioning model. **b)** Freezing rate of during the recall phase, n = 5 mice, mean +/- S.E.M. **c)** Engram cells (tdTomato+) in BLA revealed by merfish, n = 7-8 mice, mean +/- S.E.M. **d)** Unbiased clustering of all neurons. **e)** Neuronal markers and cell type annotations resolved in spatial. **f)** Unbiased clustering of neurons within BLA. **g)** Marker genes of BLA neuronal subtypes. **h)** Neuronal markers and cell type annotations of BLA. **i)** Fear memory induced gene expression in excitatory engram neurons of BLA. **j)** Fear memory induced gene expression in inhibitory engram neurons of BLA.

## A spatially resolved ensemble of engram cells

To visualize the gene expression patterns of sparsely distributed engram cells, we performed spatial transcriptomic analyses with single-molecule resolution^17^, which allowed us to study TRAPed tdT+ ‘engram’ cells *in situ* (Fig. 1**c**, Extended Data Fig. 1**a**,**b**). Fear memory consolidation in ‘FR’ mice resulted in more tdT+ engram neurons than in NF mice, especially in the BLA, paraventricular nucleus (PVT), ventral posterior complex of thalamus (VP) and zona incerta (ZI) (Extended Data Fig. 1**c**–**i**). The slice-based analysis we used not only provides spatial information, but also preserves the tissue’s native cellular architecture and avoids dissociation bias. With a customized panel of 158 genes derived from scRNAseq data (see below), we observed eight major classes of cells from over one million cells (Extended Data Fig. 1**j**,**k**), including 496,660 neurons that formed at least 17 types (Fig. 1**d**,**e**) and seven major non-neuronal cell types (Extended Data Fig. 1**j**,**k**). Consistent with a previous study^18^, neurons accounted for 48.5% of the cells.

**Extended Data Figure 1.**
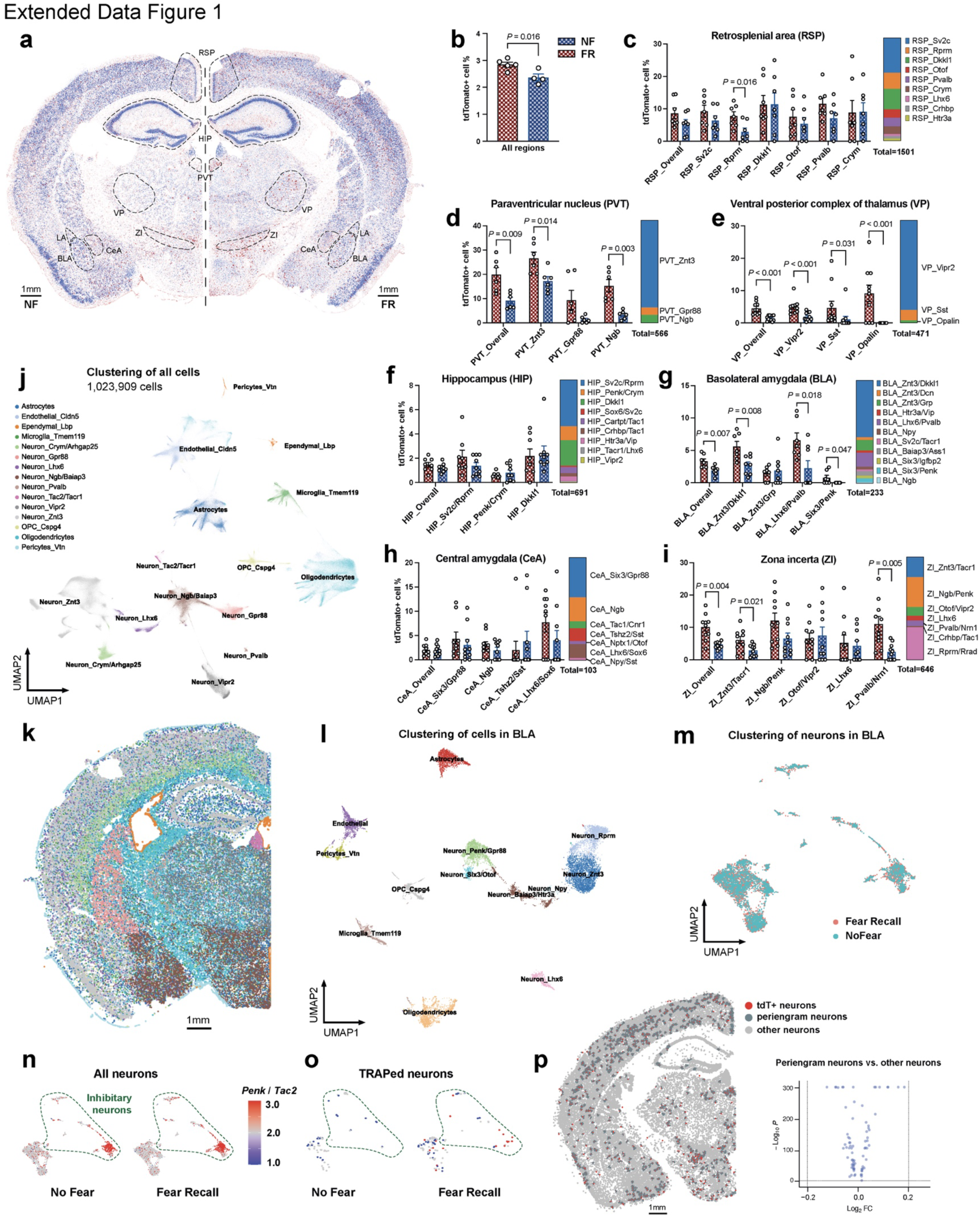
**a)** Engram cells (tdTomato+) revealed by merfish. **b-i)** Quantification of tdTomato+ neurons in all regions (**b**), retrosplenial area (RSP, **c**), paraventricular nucleus (PVT, **d**), ventral posterior complex of thalamus (VP, **e**), hippocampus (HIP, **f**), basolateral amygdala (BLA, **g**), central amygdala (CeA, **h**), and zona incerta (ZI, **i**) **j)** Major cell types with annotations resolved in a UMAP **k)** Unbiased clustering of all cells resolved *in situ*. **l)** Unbiased clustering of all cells from BLA. **m)** BLA neurons grouped by FR and NF conditions. **n)** *Penk* to *Tac2* ratio of all neurons in BLA. **o)** *Penk* to *Tac2* ratio of TRAPed neurons in BLA. **p)** Differentially gene expression analysis of Periengram neurons (neurons within a radius of 30um to engram neurons) other other neurons.

Within the BLA we identified astrocytes, microglia, oligodendrocytes, OPCs, endothelial cells, pericytes (Extended Data Fig. 1**l**) and eleven types of neurons (three excitatory and eight inhibitory) (Fig. 1**f**–**h**). The neuron types express distinctive marker genes, including *Dkkl1, Dcn, Grp* for excitatory neurons and *Vip, Pvalb, Npy, Tacr1, Baiap3, Six3, Penk* and *Ngb* for inhibitory neurons (Fig. 1**f**–**h**). Differentially activated tdT+ neurons in the FR condition presumably correspond to engram cells that are part of a persistent memory signature since we are instituting the memory recall two weeks after training and are analyzing gene expression after another nine days. However, the handling of the mice in the three control conditions, especially during the NF condition, may also activate gene expression that is unrelated to memory. Because of this circumstance and possibly due to non-specific background activation, even in the control conditions some tdT+ cells are detected. Therefore, we computed differentially expressed genes (DEGs) between the tdT+ cells in the NF and FR conditions. Since these DEGs were monitored nine days after memory recall, the DEGs likely embody genes whose expression is induced during the recall as a function of the previous fear memory training and are persistently expressed after being induced. Among 9,767 neurons in the BLA, 159 (or 3.31 %) tdT+ neurons were identified in the FR condition vs. 74 (or 1.91 %) neurons in the NF condition (Fig. 1**c**). Differential gene expression analysis in inhibitory neurons of FR engram neurons identified genes associated with synaptic vesicles (*Sv2c*) that were upregulated in the FR over the NF condition, while tachykinin 2 (*Tac2*) was downregulated. Early response genes (*Dusp1*, *Fos*) were upregulated among the excitatory neurons, whereas the neuropeptide gene *Penk* was induced in both excitatory and inhibitory neurons (Fig. 1**i**,**j**). Among tdT+ neurons, FR enriched the expression of neuropeptide *Penk* over *Tac2* more than NF, but not in total neurons (Extended Data Fig. 1**n**,**o**). In agreement with the notion that engram neurons are heterogeneously and broadly spatially distributed^19–21^, we did not observe major differences between neurons within close proximity to tdT+ neurons versus more distal neurons. (Extended Data Fig. 1**p**)

## Identification of the memory engram in the BLA and associated genes

To study engram cells in depth, we used full-length deep single-cell RNAseq experiments^22^ with an average transcript detection of 9,144 gene/neuron. We analyzed the transcriptome of 6,362 cells of the BLA, which allowed identification of all major cell types, including neurons (*Rbfox3*+), astrocytes (*Slc1a3*+), microglia (MG, *Ctss*+), oligodendrocytes (Oligo, *Plp1*+), oligodendrocyte progenitor cells (OPC, *Cspg4*+), endothelial cells (Endo, *Cldn5*+), and ependymal cells (Ependy, *Kcnj13*+)(Fig. 2**a**,**b**, Extended Data Fig. 2**a**). The relative abundance of cell types was conserved among training conditions (Extended Data Fig. 2**b**), suggesting that long-term fear memory formation does not alter the cellular architecture of the BLA.

**Fig. 2.**
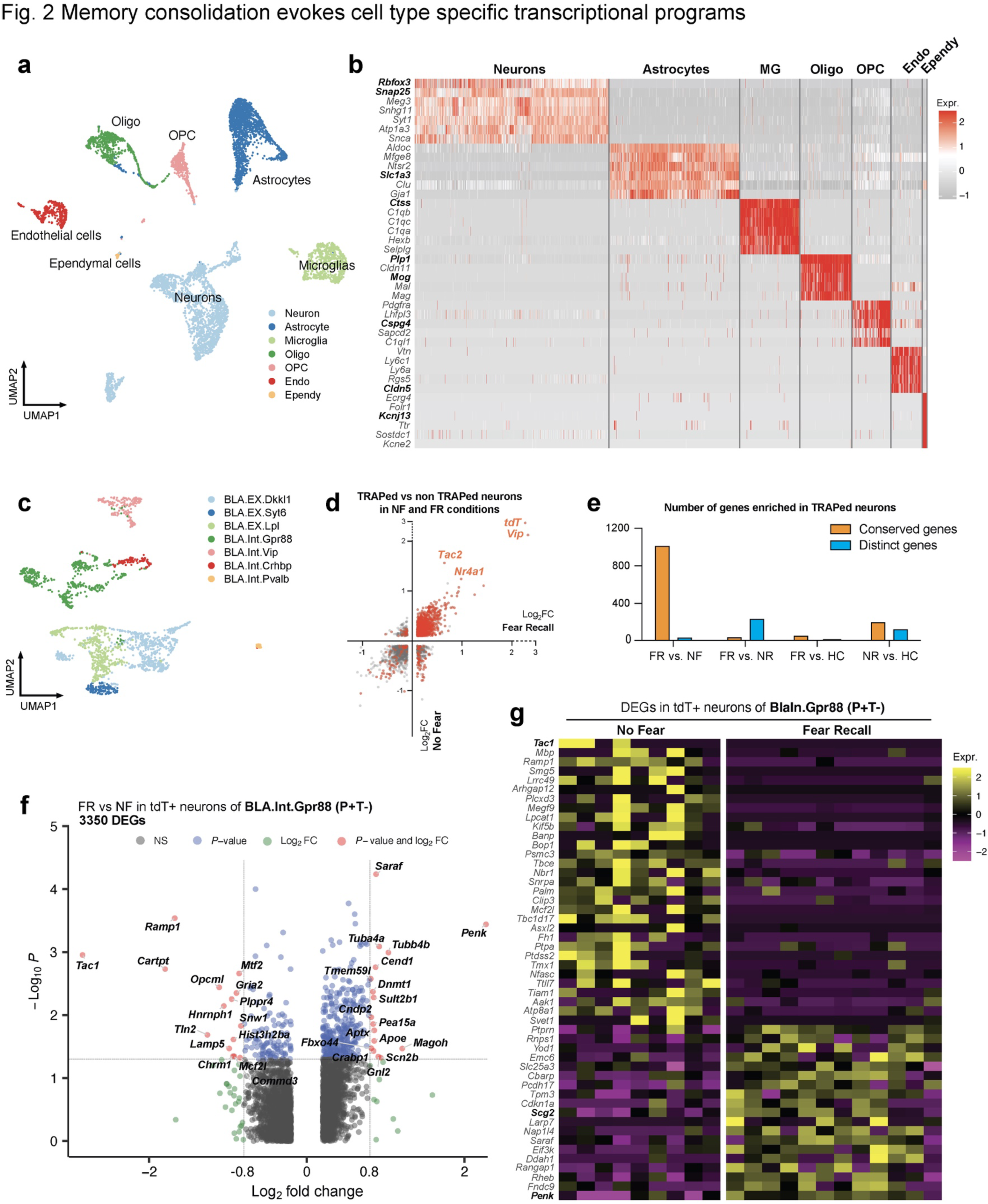
Memory consolidation evokes cell type specific transcriptional programs. **a)** Clustering of all cells in BLA, using Smartseq3 sequencing. **b)** Distinct markers for each cluster of neurons. **c)** Clustering of BLA neurons, detecting 9144 genes/cell in median. **d)** DEGs of TRAPed neurons over non TRAPed neurons in FR condition (X axis) and in NF condition (Y axis), red denotes significant DEGs (*P* < 0.05, Mann Whitney Wilcoxon test). **e)** Quantification of genes enriched in TRAPed neurons, FR and NF are mostly conserved, while FR and NR are mostly distinct. **f)** Volcano plot shows DEGs of FR over NF of TRAPed BLA.Int.Gpr88 neurons, a type of P+T- neurons. **g)** DEGs of FR over NF of TRAPed BlaIn.Gpr88 neurons, a type of P+T- neurons.

**Extended Data Figure 2.**
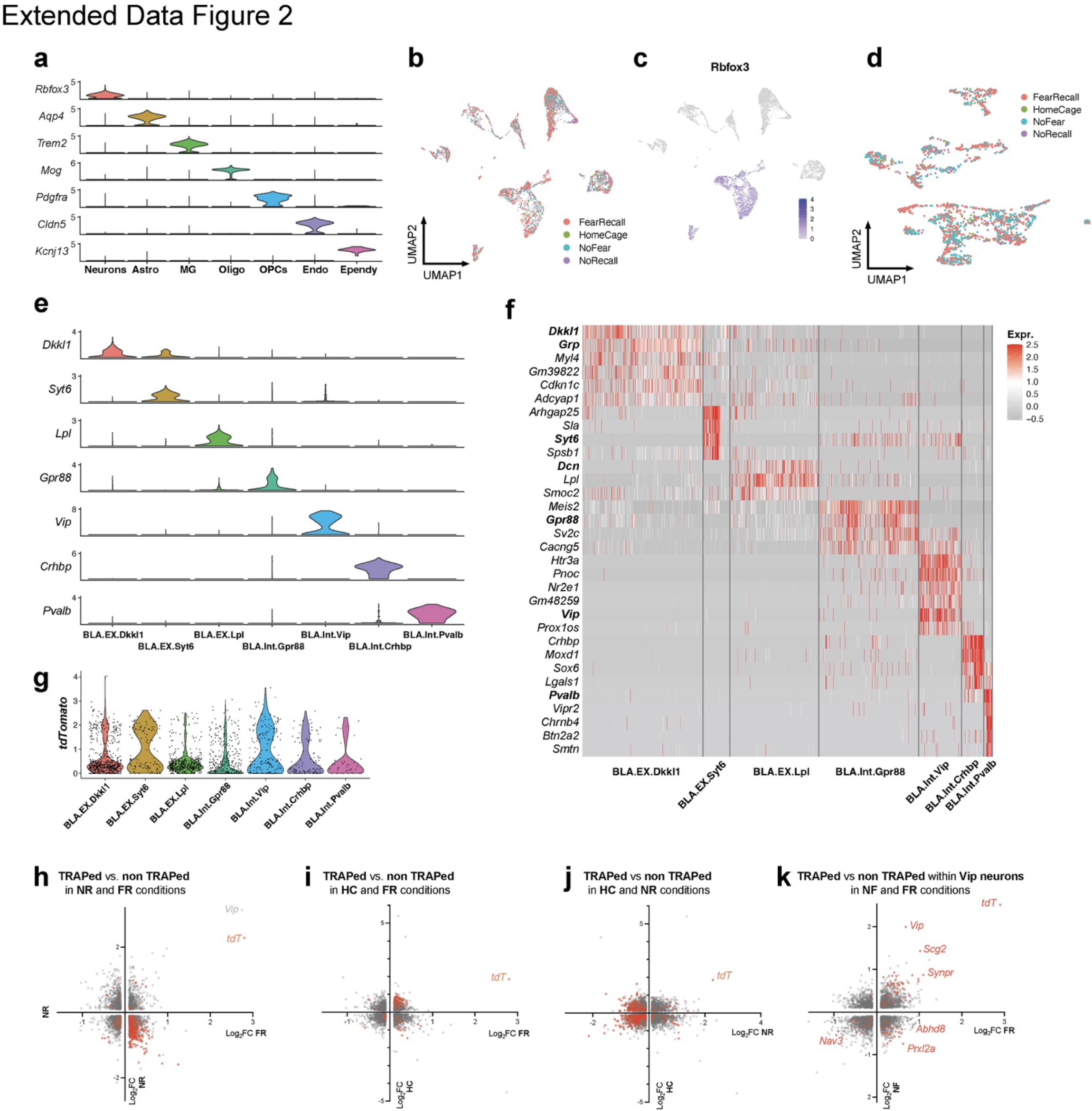
**a)** Distinct markers for each cluster of BLA cells. **b)** BLA cell clustering colored by training conditions. **c)** *Rbfox3* expression in BLA cells. **d)** BLA neurons clustering colored by training conditions. **e)** Distinct markers for each cluster of BLA neurons. **f)** Heatmap of top marker genes of neuronal clusters **g)** tdTomato expression in each neuron cluster. **h**-**k**) DEGs of TRAPed neurons over non TRAPed neurons, red denotes significant DEGs.

Subclustering of 2,144 neurons (456 of which were tdT+) revealed seven subtypes (Fig. 2**c**,Extended Data Fig. 2**e**,**f**) that express multiple distinctive marker genes. These subtypes were consistently observed in the four training conditions (Extended Data Fig. 2**d**) and validated by the spatial transcriptomic data (Extended Data Fig. 2**e**,**f**, Fig. 1**f**,**g**). All subtypes contained tdT+ cells, suggesting that all subtypes were activated during recall (Extended Data Fig. 2**g**). We then analyzed which genes characterize tdT+ cells. Besides tdT, genes encoding neuropeptides (e.g., vasoactive intestinal peptide (*Vip*) and tachykinin 2 (*Tac2*)) and the immediate early gene *Nr4a1* were enriched in tdT+ neurons. These genes were consistently observed in both the FR and NF conditions (Fig. 2**d**), but not the HC and NR conditions (Fig. 2**e**, Extended Data Fig. 2**h**–**k**), suggesting that the salient experience of placing the mice into the fear conditioning chamber in the NF condition is sufficient to induce a long-lasting gene expression change lasting. Interestingly, in line with our observation that *Vip* is the most prominently induced gene in tdT+ neurons it has been reported that *Vip* interneurons are activated by salient cues in the BLA and that such activation is required for learning.^23^ However, given that *Vip* was induced even in the NF condition, it clearly is not an engram gene.

## Memory-associated gene expression

Three of the seven types of BLA neurons are glutamatergic (BLA.EX.Dkkl1/Syt6/Lpl) and four are GABAergic (BLA.Int.Gpr88/Vip/Crhbp/Pvalb) (Extended Data Fig. 3**a**). Interestingly, the FR condition recruited a significantly higher number of tdT+ neurons than the NF condition within the BLA.Int.Gpr88 population (Extended Data Fig. 3**b**), which is marked by *Gpr88*, synaptic vesicle glycoprotein 2C (*Sv2c*), and calcium voltage-gated channel auxiliary subunit gamma 5 (*Cacng5*) (Extended Data Fig. 2**f**).

**Fig. 3.**
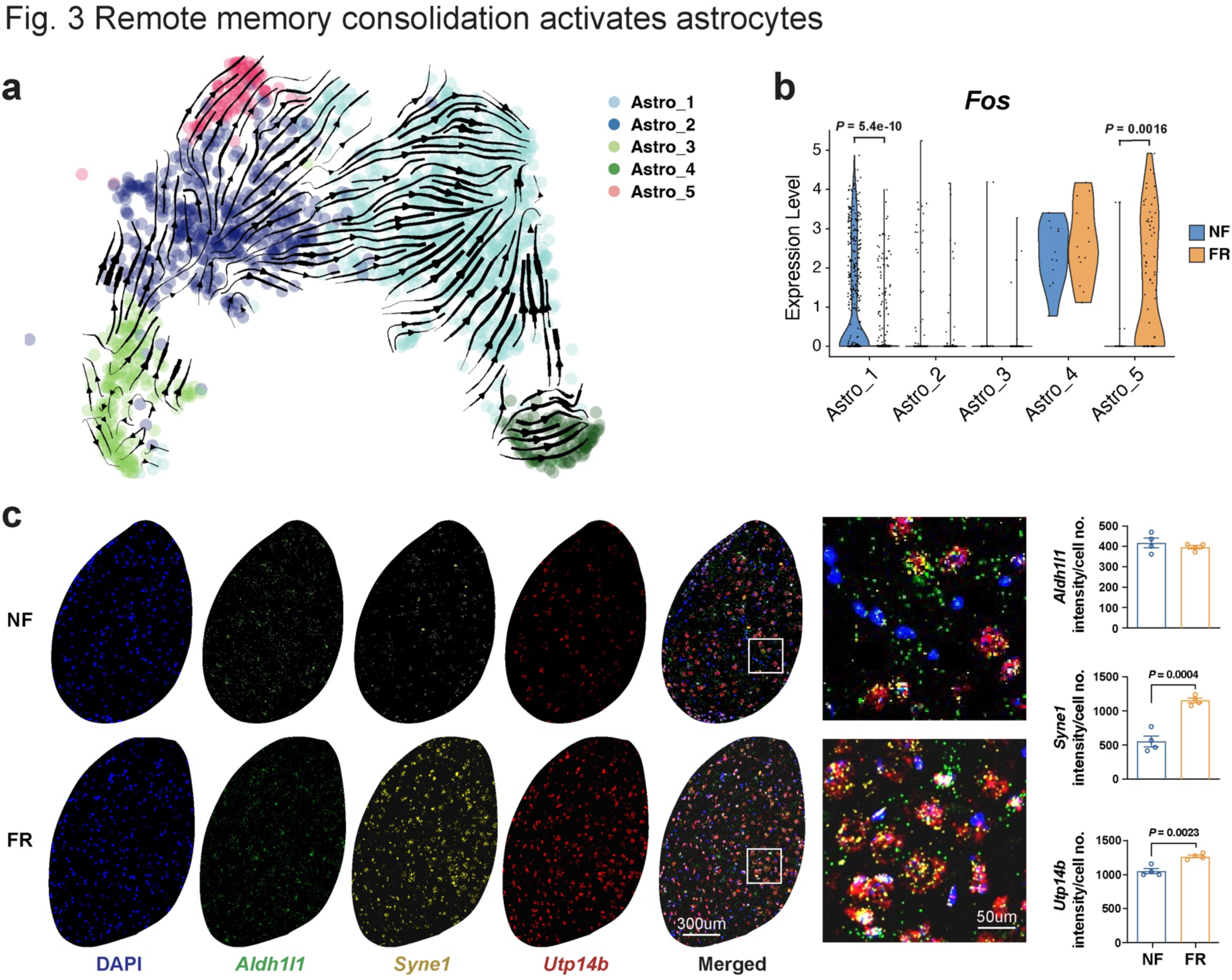
Remote memory consolidation activates astrocytes. **a)** Cellular trajectory estimation of BLA astrocytes, based on RNA maturation. **b)** *Fos* expression of FR and NF astrocyte **c)** RNAscope *in situ* stanning of *Aldh1l1*, *Syne1*, and *Utp14b* in BLA of FN and FR.

**Extended Data Figure 3.**
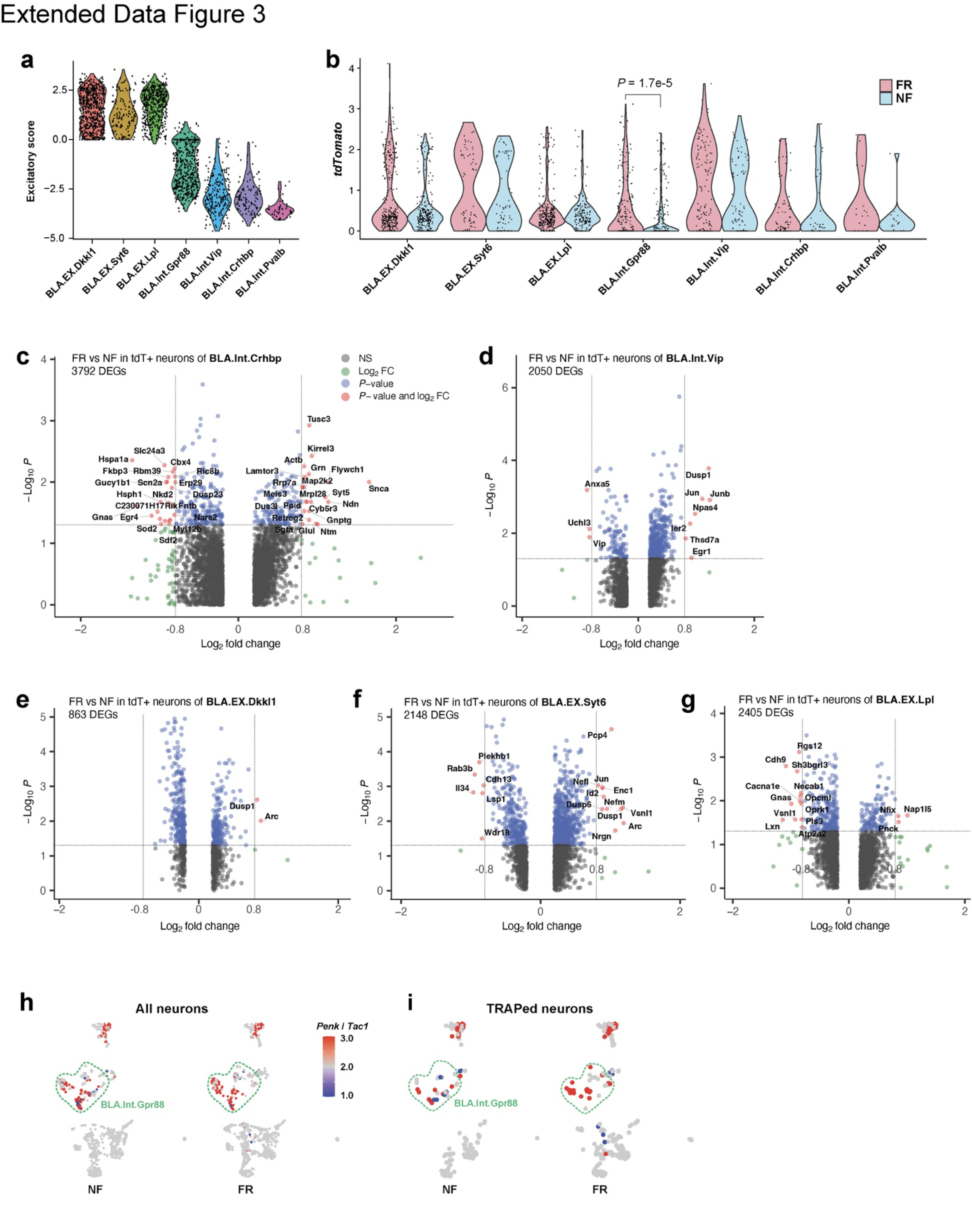
**a)** Excitatory score of BLA neurons, calculated by Scl17a7 – Gad1. **b)** tdTomato expression in each neuron cluster, splited by training conditions, two-tailed student T-test. **c**-**g)** DEGs of FR over NF of TRAPed BLA.Int.Crhbp (**c**), BLA.Int.Vip (**d**), BLA.EX. Dkkl1 (**e**), BLA.EX.Syt6 (**f**), and BLA.EX.Lpl (**g**) neurons **h)** Penk to Tac1 ratio of all neurons in BLA. **i)** Penk to Tac1 ratio of TRAPed neurons in BLA.

To identify transcriptional changes specifically induced by remote memory recall in engram neurons, we screened for DEGs in TRAPed tdT+ neurons of FR versus NF mice. Single-cell resolution enables a comparison of neurons of the same type and full-length mRNA sequencing provides in-depth identification of genes that are specifically associated with memory consolidation and recall. Strict criteria were applied to remove non-specific DEGs. First, DEGs that were also differentially expressed between non-TRAPed cells in FR versus NF mice were removed, which minimized the impact of basal activation. Second, only DEGs that were differentially expressed when FR cells are compared to NR and HC controls were included, ensuring that DEGs are not just a consequence of a fear experience. Lastly, each DEG had to be expressed in at least one-quarter of cells and with at least a 1.75-fold change. These stringent criteria identified 107 ‘remote-memory-associated DEGs’ in six types of neurons (Fig. 2**f**, Extended Data Fig. 3**c**–**g**).

Inhibitory neurons in the BLA have been shown to regulate fear memory consolidation^24–26^ in a cell type specific manner.^23, 27^ Interestingly, we found that the GABAergic inhibitory neurons BLA.Int.Gpr88 and BLA.Int.Crhbp showed more regulation than the other neurons (Fig. 2**f**, Extended Data Fig. 3**c**–**g**), suggesting that inhibitory neurons in BLA are more actively involved in memory consolidation. Strikingly, the largest effect of remote memory recall was observed in two neuropeptide genes that different from those detected in salience-activated gene expression changes: *Tac1*, whose expression was suppressed over 6-fold, and *Penk*, whose expression was increased more than 4-fold in BLA.Int.Gpr88 neurons (P+T- neurons) (Fig. 2**f**). As a result, tdT+ engram neurons in the FR condition showed a much higher *Penk* to *Tac1* ratio than tdT+ neurons in the NF condition in BLA.Int.Gpr88 neurons (Extended Data Fig. 3**h**,**i**). In addition, we found a strong enrichment in genes involved in mitogen-activated protein kinase pathways (*Dusp1*, *Dups6*, *Nefl*, *Lamtor3*, *Jun*, *Junb*, and *Map2k2*) (Fig. 2**f**, Extended Data Fig. 3**c**–**g**), consistent with the implication of MAPK pathways in memory consolidation in a variety of learning paradigms^28–32^, including fear memory consolidation in the amygdala^33^. Genes related to signaling in general, in particular brain-derived neurotrophic factor (BDNF) signaling (*Egr1*, *Vsnl1*, *Dusp1*, *Hnrnph1*, *Id2*, *Ramp1*, *Ier2*, *Hspa1a*) (Fig. 2**f**, Extended Data Fig. 3**c**–**g**), were also found to be differentially regulated by fear memory. In the amygdala, BDNF signaling was reported to be essential for fear memory consolidation^34, 35^, fear memory extinction^36^, episodic memory formation^37^, and long-term potentiation^38^. Interestingly, BDNF and MAPK were shown to relay signaling cascades and enhance stress-induced contextual fear memory^39^.

More than half of the DEGs associated with remote memory also have links to neuronal disorders such as dementia, mental retardation, epilepsy, schizophrenia, and Charcot-Marie-Tooth Disease type I and II. This indicates a potential correlation between the functional role of these genes in regulating remote memory and their involvement in the development of neurological disorders.

Because inhibitory neurons in BLA showed dramatic engram specific gene regulation, we further subclustered inhibitory neurons into five subtypes BlaIn.Sst/Vip/Gpr88/Calm1/Pvalb (Extended Data Fig. 4**a**–**e**). Differential analysis of these TRAPed tdT+ inhibitory neuron subtypes between FR and NF uncovered 159 genes that were associated with memory consolidation (Fig. 2**g**, Extended Data Fig. 4**f**–**h**). Transcription factor enrichment analysis^30^ of the FR induced genes revealed a strong enrichment of target genes of cAMP responsive element binding protein 1 (CREB) (Extended Data Fig. 4**j**). The CREB signaling pathway is widely implicated in long-term memory consolidation.^6–8^

**Fig. 4.**
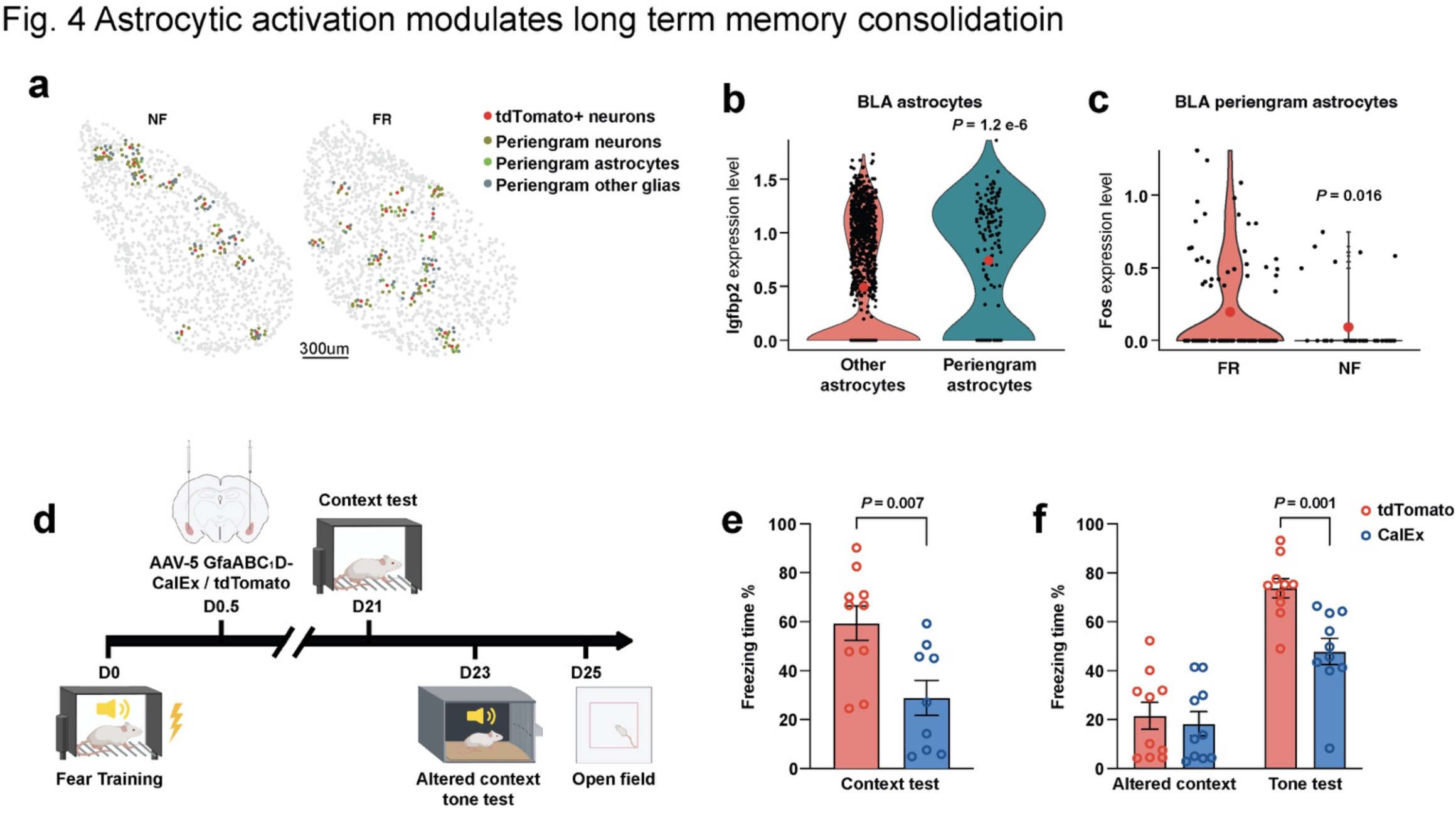
Astrocytic activation modulates long erm memory consolidation. **a)** Spatial resolved cells surrounding tdT+ neurons. **b)** *Igfbp2* expression is enriched in astrocytes surrounding tdT+ neurons (Mann Whitney Wilcoxon test). **c)** *Fos* expression in induced in FR condition than NF condition, among periengram astrocytes (Mann Whitney Wilcoxon test). **d)** Scheme, adeno associated virus conveying GfaABC1D-mCherry-CalEx (or GfaABC1D- mCherry) were bilaterally injected to BLA C57B/6 mice, one day after fear conditioning training. Mice were subjected to context test, altered context tone test, and open field test at time indicated in the scheme. **e)** Mice with CalEx showed reduced freezing than mCherry control group in context test, n = 9-10 mice, mean +/- S.E.M, two tailed student T-test. **f)** Mice with CalEx showed comparable freezing in altered context but reduced freezing in tone test, n = 10 mice, mean +/- S.E.M, two tailed student T-test.

**Extended Data Figure 4.**
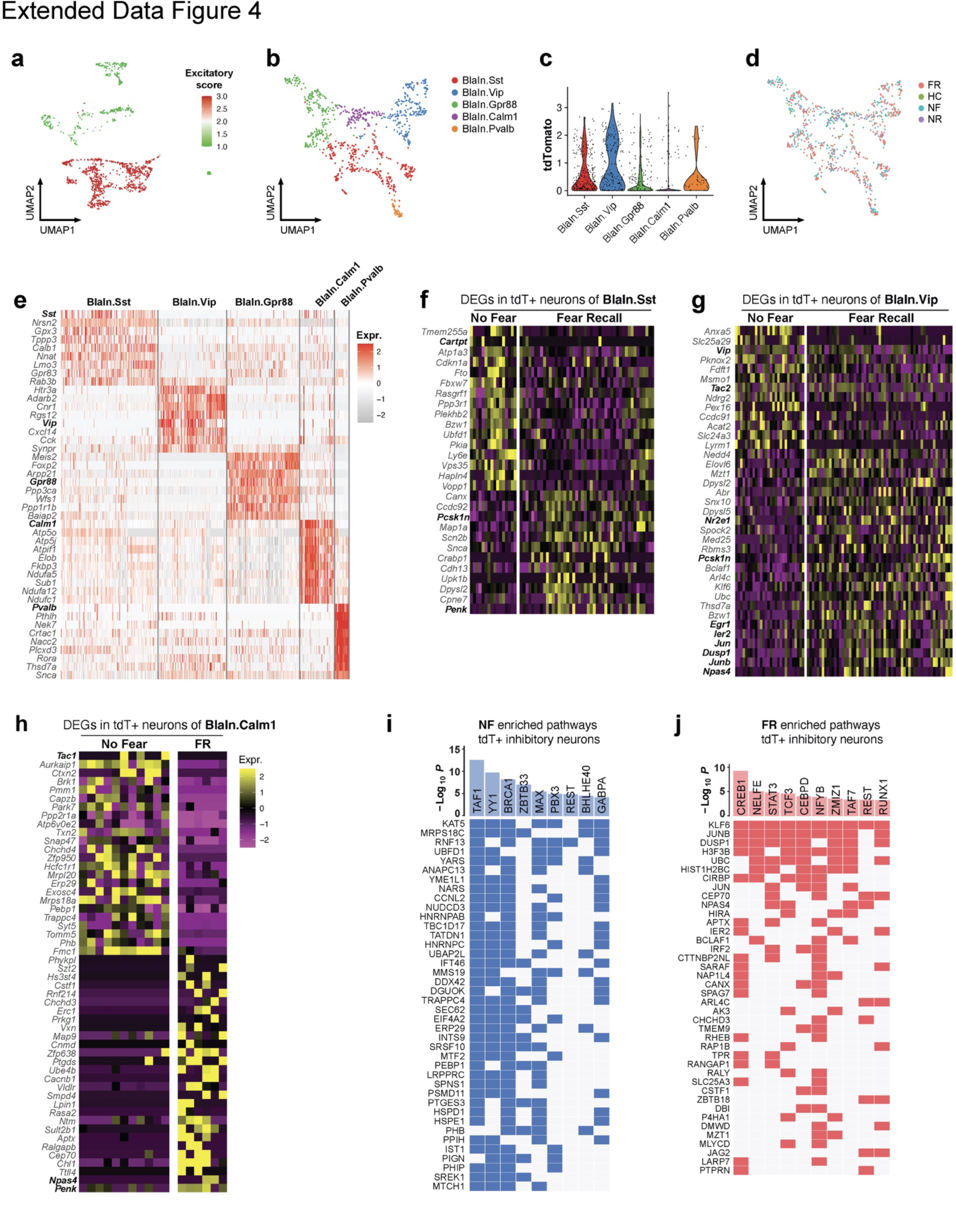
**a)** Excitatory score of BLA neurons, calculated by Scl17a7 – Gad1. **b)** BLA inhibitory neuron clustering. **c)** tdTomato expression in each inhibitory neuron cluster. **d)** BLA inhibitory neuron clustering colored by training conditions. **e)** Heatmap of top marker genes of inhibitory neuronal clusters **f**-**h**) DEGs (FR over NF, TRAPed) of BlaIn.Sst (**f**), BlaIn.Vip (**g**), and BlaIn.Calm1 (**h**). **i**-**j**) Transcription factor enrichment analysis of NF induced genes (**i**) or FR induced genes (**j**).

A set of immediate-early genes was also regulated by fear memory consolidation, including the early immediate early response 2 (*Ier2*), early growth response 1 (*Egr1*), *Jun*, *Junb*, dual specificity phosphatase 1 (*Dusp1*), and neuronal PAS domain protein 4 (*Npas4*) genes (Fig. 2**g**, Extended Data Fig. 4**f**–**h**). In particular, *Egr1* was reported to be required in lateral amygdala for long-term fear memory consolidation without impairing acquisition or short-term memory.^40^ *Npas4* is a Ca^2+^ influx dependent gene that regulates synapse development in inhibitory neurons by regulating activity-dependent genes^41^, marks a subset of fear induced engram neurons in parallel of Fos engrams^42^, is required for both short-term and long-term contextual fear memory^43^. Previous work in hippocampus showed *Penk*, *Dusp1*, *CREB*, *Npas4* are involved in fear memory^44^.

Interestingly, we found a host of genes associated with neuropeptides were regulated during fear memory consolidation: secretogranin 2 (*Scg2*)^45^ and *Penk* were upregulated while tachykinin 1 (*Tac1*) were down regulated among BlaIn.Gpr88 neurons (P+T- neurons) (Fig. 2**g**). Similarly, *Penk* was upregulated in both BlaIn.Sst and BlaIn.Calm1 neurons, in which cocaine- and amphetamine-regulated transcript protein (CART, encoded by *Cartpt*) and *Tac1* down regulated respectively (Extended Data Fig. 4**f**,**h**). In Vip neurons, tachykinin 2 (*Tac2*) and vasoactive intestinal peptide (*Vip*) were down regulated (Extended Data Fig. 4**g**). ProSAAS (*Pcsk1n*, a neuroendocrine peptide precursor) was upregulated in both BlaIn.Sst and BlaIn.Vip neurons (Extended Data Fig. 4**f**,**g**), *Pcak1n* was reported to inhibit prohormone processing,^46^ and be required for fear memory.^47^ This data suggests engram neurons switched the production of neuropeptides during memory consolidation.

## Astrocyte remodeling in remote memory consolidation

Neuron–glia interactions are thought to play an essential role in memory consolidation.^48–50^ Moreover, astrocytes respond to neuronal activity with neuronal activity-dependent sharp tuning^51^. Given the fact that *Nts* is induced during memory formation in engram neurons and that neurotensin encoded by *Nts* can directly activate astrocytes by stimulating the GPCR *Ntsr2*, we asked whether memory-induced persistent gene expression changes also exist in astrocytes. Among non-neuronal cells, only astrocytes exhibited consistent transcriptional changes associated with remote memory consolidation. Unbiased clustering of 1,637 astrocytes identified five cell states that might be considered astrocyte subtypes (Astro_1-5) (Extended Data Fig. 5**a**–**c**). Cellular trajectory analyses based on RNA dynamics^52^ and gene expression patterns^53^ suggested an astrocytic cellular pathway from Astro_3 to Astro_2 to Astro_1 to Astro_4, with a split from Astro_2 to Asro_5 (Fig. 3**a**, Extended Data Fig. 5**d**).

**Fig. 5.**
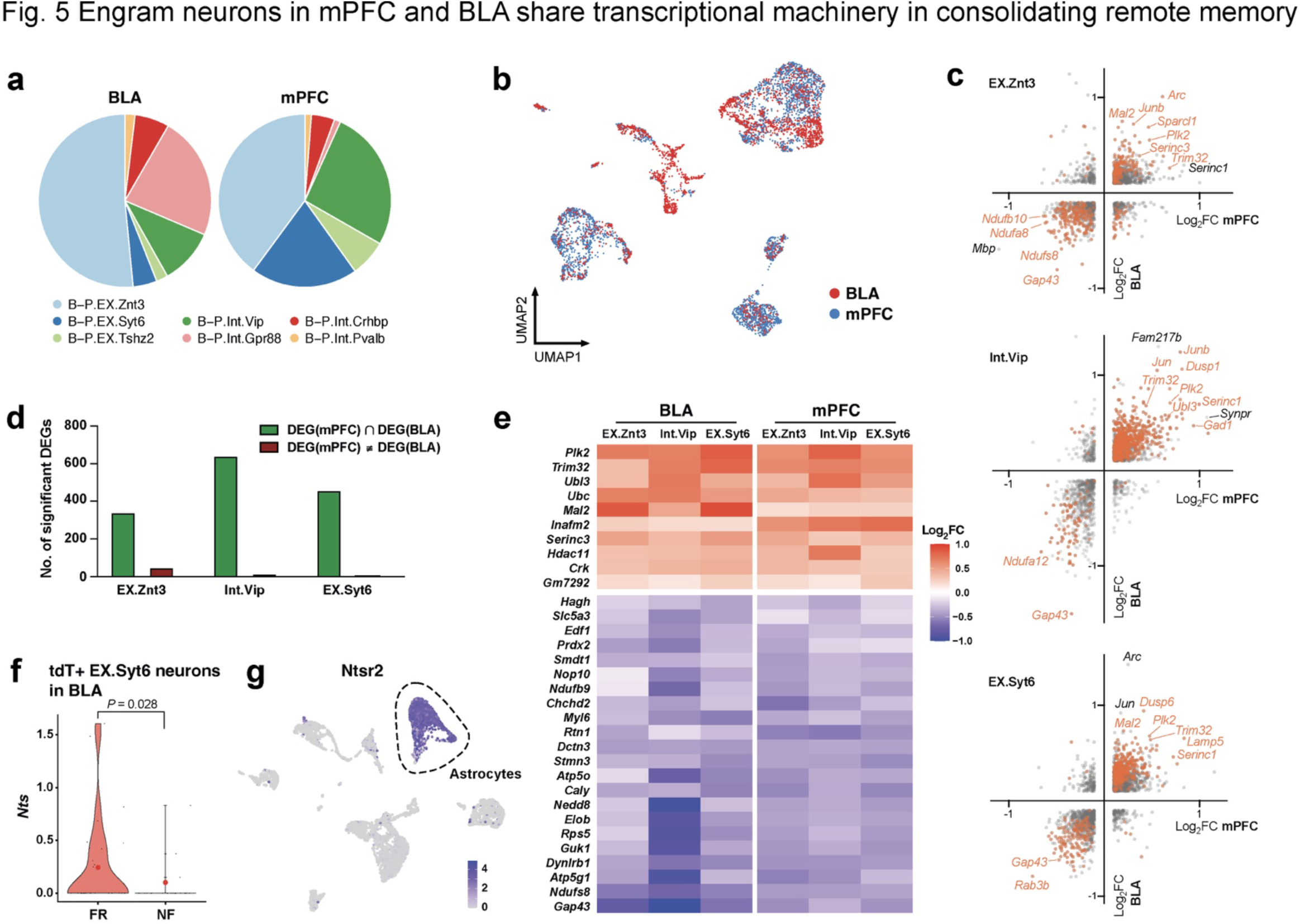
Engram neurons in mPFC and BLA share transcriptional machinery in consolidating remote memory a) Cellular composition of BLA and mPFC. b) Integrated clustering of BLA and mPFC neurons, colored by regions. c) DEGs of TRAPed cells of EX.Znt3 (top), Int.Vip (middle), and EXT.Syt6 (bottom). X axis is fold change of FR over NF in BLA, Y axis is fold change of mPFC. Orange denotes significant DEGs (*P* <0.05, Mann Whitney Wilcoxon test). d) Quantification of significant DEGs in neuron clusters 1-3. e) DEGs (FR over NF, TRAPed cells) from BLA and mPFC among B-P.EX.Znt3, B-P.Int.Vip, and B-P.EX.Syt6 neurons. f) *Nts* expression in tdT+ B-P.EX.Syt6 neurons from BLA. g) *Ntsr2* expression in all cell from BLA, *Ntsr2* expression is highly enriched in astrocytes.

**Extended Data Figure 5.**
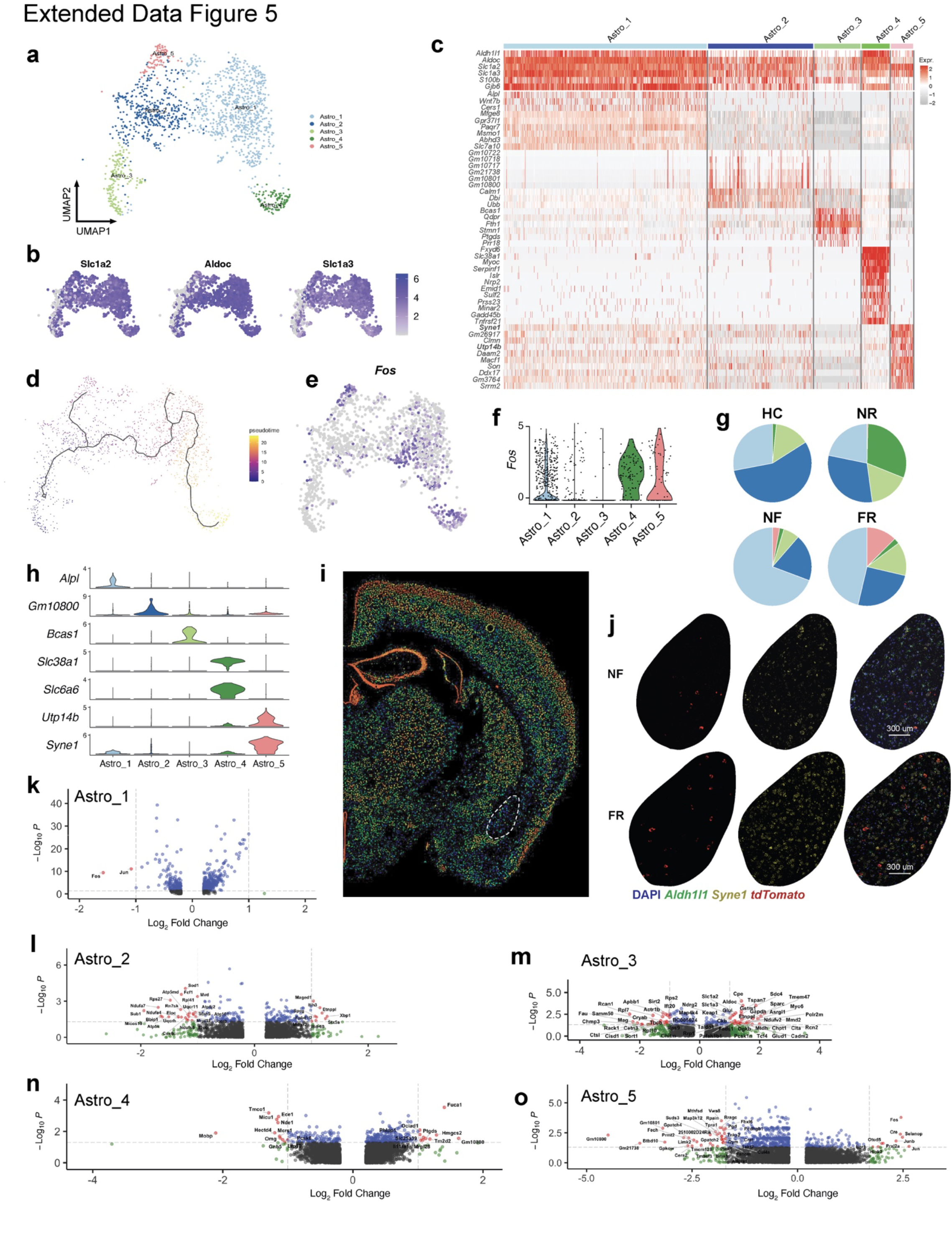
**a)** Cluster of astrocytes from BLA. **b)** Expression level of astrocyte pan markers (*Slc1a2*, *Aldoc*, and *Slc1a3*). **c)** Heatmap of top marker genes of BLA astrocyte clusters. **d)** Cellular trajectory estimation of BLA astrocytes, based on gene expression. **e)** *Fos* expression of BLA astrocytes. **f)** *Fos* expression of astrocyte clusters **g)** Astrocyte composition separated by training conditions. **h)** Distinct markers for each astrocyte cluster from BLA. **i)** Syne1 expression data, retrieved from Allen Atlas. **j)** RNAscope *in situ* stanning of *Syne1* and *tdTomato* in BLA of NF and FR conditions. **k-o)** DEGs of FR vs. FN in Astro_1 – 5.

Astrocyte activation has been increasingly recognized as important in multiple types of memory^54^. Astro_4 showed high expression of *Fos,* suggesting the final astrocyte cell state is an active state (Extended Data Fig. 5**e**,**f**). Of note, Astro_4 cells also express the GABA transporter *Slc6a6*^55^ and the glutamine transporter *Slc38a1*^56^ (Extended Data Fig. 5**c**,**h**), suggesting functional roles in regulating neurotransmitter levels. In addition, Astro_1 and Astro_5 also showed activation (Extended Data Fig. 5**e**,**f**). Remarkably, memory consolidation promoted the split from Astro_2 to Astro_5 and reduced the Astro_1 proportion comparing to NF condition (Fig. 3**a**, Extended Data Fig. 5**d**,**g**). Astro_1 cells are significantly less active in the FR than in the NF condition, whereas Astro_5 are more active, suggesting that memory consolidation shifted the active astrocytes from Astro_1 to Astro_5 (Fig. 3**b**), which express *Utp14b*, *Syne1, Son, Macf1* and other marker genes (Extended Data Fig. 5**c**,**h**).

An astrocyte subtype with markers of *SON*, *MACF1* and *SYNE1* was recently identified in the human anterior cingulate cortex^57^. *Utp14b* was found to be upregulated in astrocytes of the neocortex in stressed mice ^58^. Humans with *Syne1* mutant are more likely to develop autism^59^ and bipolar disorder^60^. The expression of *Syne1* is relatively low in BLA under basal conditions (Fig. 3**c**, Extended Data Fig. 5**i**,**j**) but was induced in the FR condition (Fig. 3**c**, Extended Data Fig. 5**j**). 133 genes were found differentially expressed in FR condition over NF. In the active state Astro_4, glutathione-independent prostaglandin D synthase (*Ptgds*) and mitochondrial glutathione transporter (*Slc25a39*) were significantly induced by FR (Extended Data Fig. 5**n**), suggesting prostaglandin is involved in memory consolidation. Meanwhile genes associated with glutamate transport (*Slc1a2* and *Slc1a3*) and glutamine synthesis (*Glul*) were upregulated in FR than NF in Astro_3 (Extended Data Fig. 5**m**).

Consistent with the scRNAseq data, we also found in the spatial transcriptomics data a subcluster of astrocytes which were induced by the FR condition (Extended Data Fig. 6**a**–**c**). This subcluster is preferentially activated in FR mice and expresses high levels of *Syne1*, *Utp14b* (Extended Data Fig. 6**d**,**e**), and *Flt1* (Extended Data Fig. 6**f**,**g**). Interestingly, *Flt1* is the receptor of vascular endothelial growth factors that is expressed in activated astrocytes^61–63^, may induce angiogenesis^61, 64^, and could facilitate synaptogenesis^65^.

**Extended Data Figure 6.**
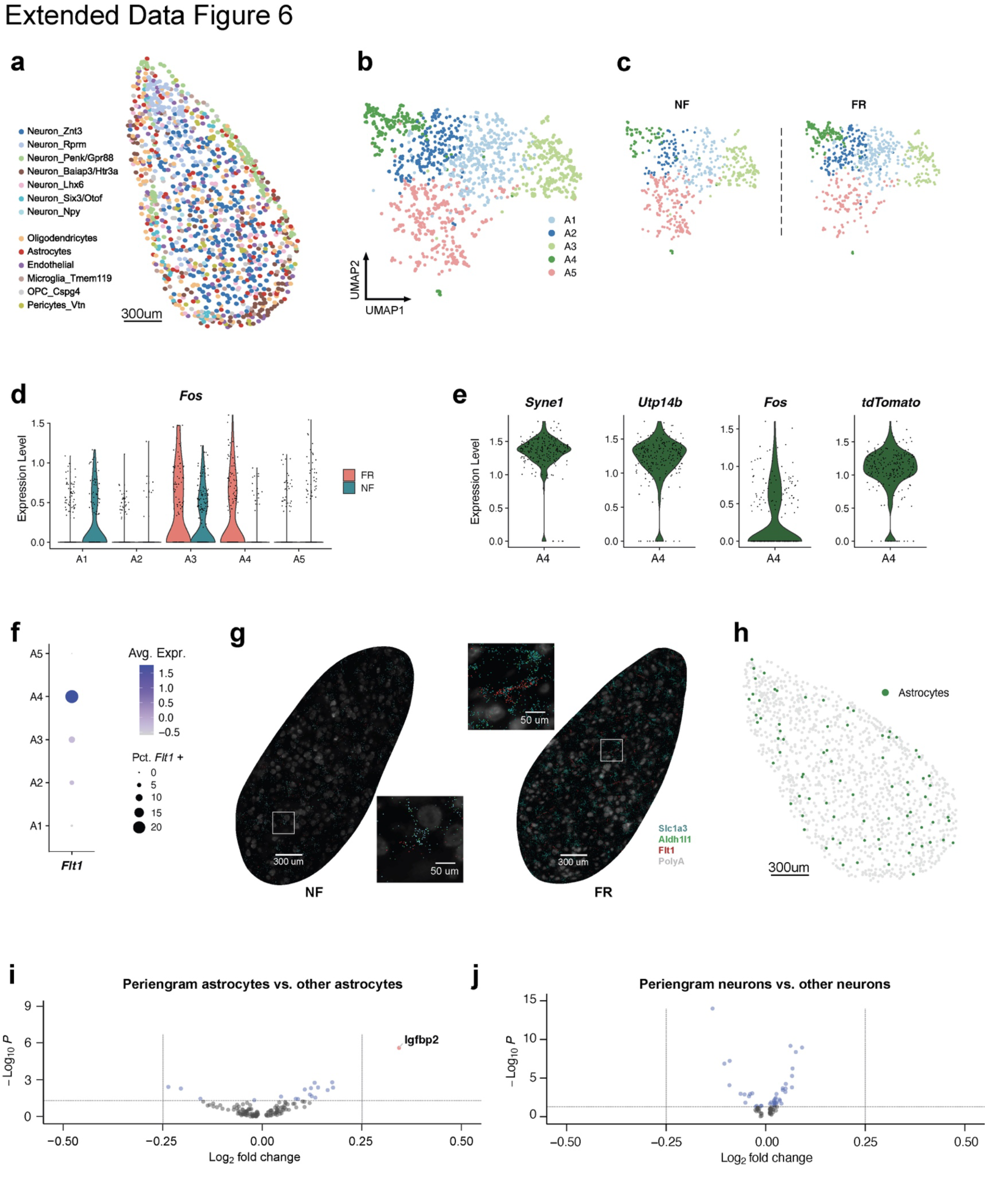
**a)** Spatial embedding of all BLA cell types from merfish data. **b)** Clustering of astrocytes in BLA from merfish data. **c)** Clustering of astrocytes in BLA from merfish data, saperated by training conditions. **d)** *Fos* expression in BLA astrocyte subtypes separated by conditions. **e)** *Syne1*, *Utp14b*, *Fos*, *tdTomato* expression in A4 astrocytes from BLA in FR, merfish data. **f)** *Flt1* expression in BLA astrocyte subtypes from merfish data. **g)** Slc1a3, Aldh1l1, and Flt1 *in situ* data from merfish. **h)** Spatial distribution of astrocytes in BLA. **i)** Volcano plots showing the genes differentially expressed in periengram astrocytes. **j)** Volcano plots showing the genes differentially expressed in periengram neurons.

## Astrocyte-to-neuron interaction in remote memory consolidation

Engram neurons are thought to be randomly distributed in the amygdala^19^ and other brain regions^20, 21^. However, some spatial structures, such as perineuronal nets, are known to be essential for these engram neurons^66, 67^. We asked whether a particular spatial cellular environment is associated with the formation of engram neurons. By analyzing the cells surrounding tdT+ neurons in the BLA (within a radius of 30 mm) (Fig. 4**a**), we found that expression of *Igfbp2* (which encodes insulin-like growth factor binding protein 2) was enriched in peri-engram astrocytes, whereas peri-engram neurons were found to be indistinguishable from other neurons (Fig. 4**b**, Extended Data Fig. 6**h**–**j**). IGFBP2, an astrocytic secreted protein, was reported to have multiple effects on neurons, including changes in synaptic transmission and excitability^68^. Interestingly, peri-engram astrocytes in the FR condition showed a higher Fos activation than in the NF condition (Fig. 4**c**). Our spatial transcriptomic data not only localized the sparse engram cells and identified the signatures of cells in close vicinity to engram cells, but also recapitulated the scRNAseq defined cellular structure and gene expression of engram, and the activation of astrocytes by memory consolidation.

To functionally assess whether activation of astrocytes contributes to memory formation, we selectively inhibited astrocyte activation in the BLA during fear memory formation. This was accomplished using expression of CalEx that removes calcium from astrocytes^69^ (Fig. 4**d**–**f**). After fear training, mice were bilaterally injected with AAVs expressing CalEx under control of the astrocyte-specific GfaABC1D promoter, using mCherry as a marker and tdTomato-only expression as a control. 21 days later, mice were subjected to contextual memory tests using first the original and then an altered context, followed by a cued fear conditioning test and open field measurements (Fig. 4**d**). We found that both contextual and cued fear conditioning memory were impaired by the suppression of astrocyte activation, whereas the response to the altered context remained unchanged and low (Fig. 4**e**,**f**). No change in the open field test was detected upon astrocyte inhibition (Extended Data Fig. 7**a**–**c**). Interestingly, a recent study showed activating astrocytes in BLA promotes fear memory formation^70^. Previously, it has been shown that activating CA1 astrocytes enhances memory allocation with increased neuronal activity in learning^71^, astrocyte activation in hippocampus is required for long-term memory^72^, and CA1 astrocyte activation is involved in encoding reward location^73^. This evidence further supports the notion that the activity of astrocytes is functionally linked to memory formation.

**Extended Data Figure 7.**
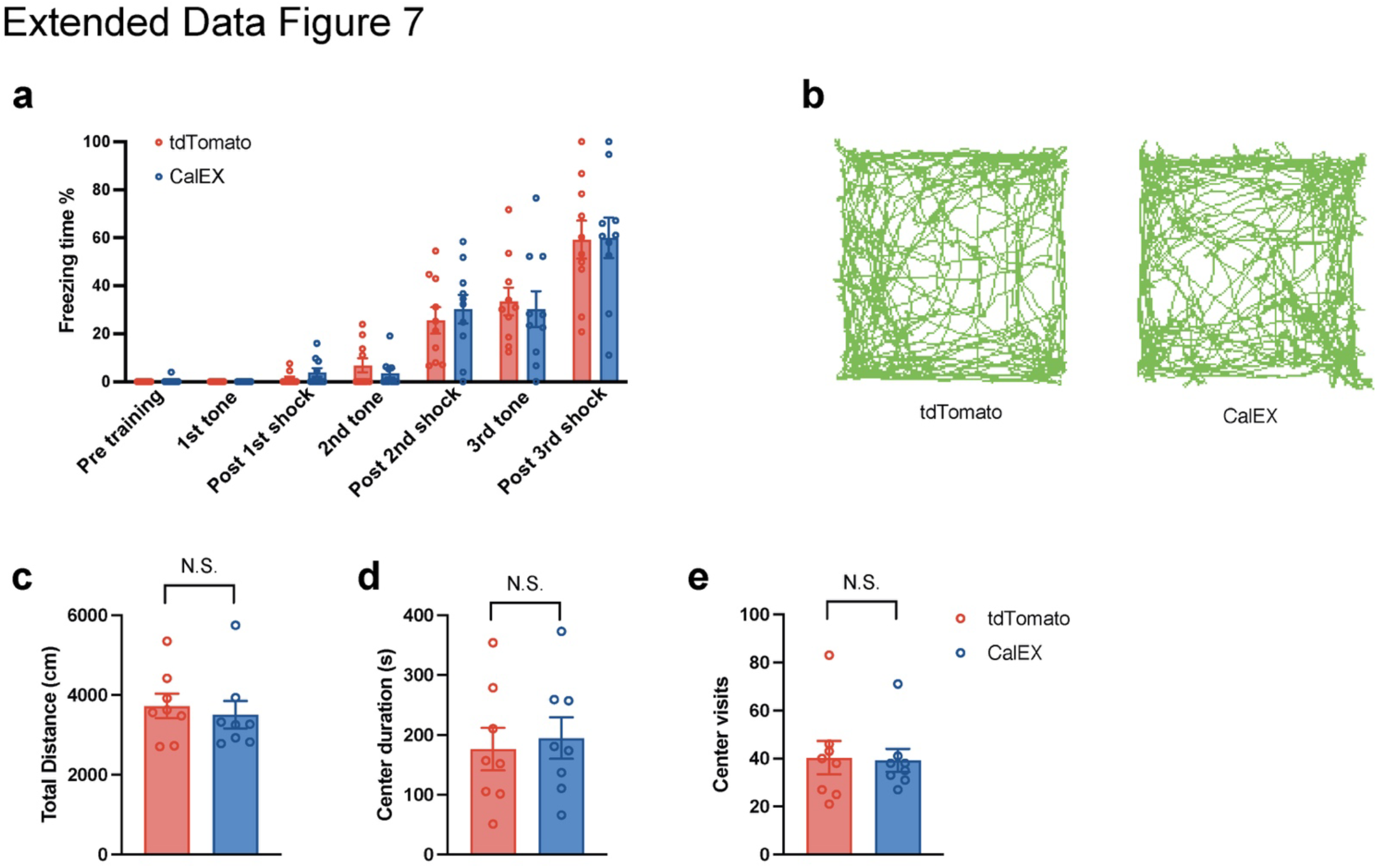
**a)** Freezing time in training, n = 8 mice, average +/- SEM. **b)** Representative tracks in open field test. **c)** Total distance, center visits, and center duration in open field test, n = 8 mice, average +/- SEM, two-tailed student T-test.

## A memory link between prefrontal cortex and amygdala

Memory formation is orchestrated by multiple connecting parts of the brain, with the prefrontal cortex and amygdala as a key axis in fear memory formation^74^. Here, we used our previously published single-cell RNAseq data from the mPFC^15^ (Extended Data Fig. 8**a**–**g**) to perform an integrated analysis of neurons from the BLA and mPFC and test whether a common gene-expression signature connects long-term memory formation in these two regions.

**Extended Data Figure 8.**
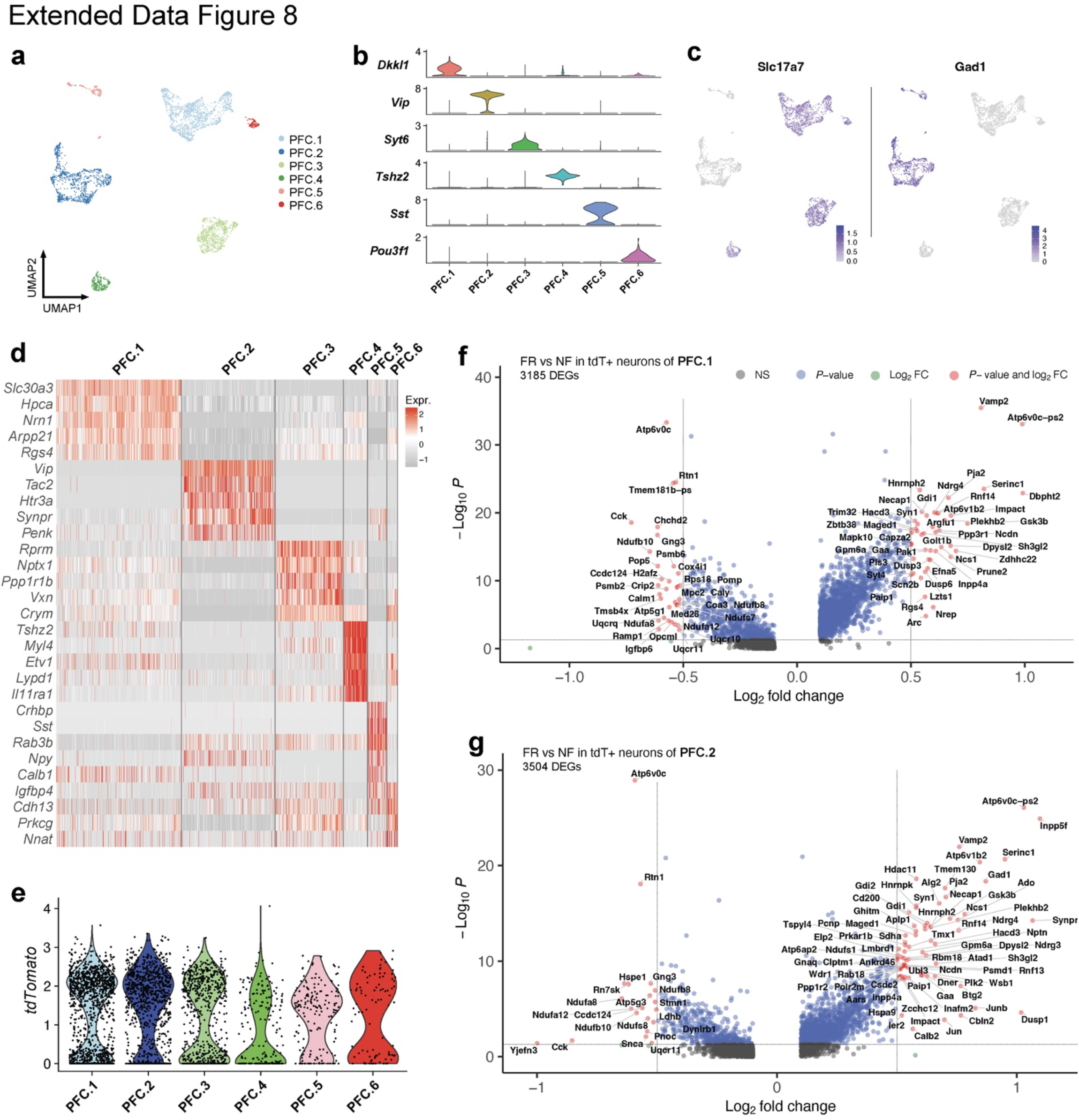
(reanalysis of scRNAseq data of mPFC neurons, Chen *et. al.*, 2020) **a)** Cluster of mPFC neurons **b)** Distinct markers for each cluster of mPFC neurons. **c)** *Slc17a7* and *Gad1* expression of mPFC neurons. **d)** Heatmap of top marker genes of mPFC neurons. **e)** tdTomato expression of mPFC neurons. **f)** DEGs of TRAPed cells from PFC.1. **g)** DEGs of TRAPed cells from PFC.2.

A set of 4,603 neurons from the mPFC and BLA were clustered into seven populations with clear markers for each cell type (Extended Data Fig. 9**a**–**d**). Interestingly, six of the seven types of neurons were found in both BLA and mPFC; only Gpr88 neurons were specific to the BLA (Fig. 5**a**,**b**, Extended Data Fig. 9**b**). EX,Znt3/Syt6/Tshz2 are excitatory neurons that express vGlut2 (*Slc17a7*), whereas the other clusters are Gad1+ inhibitory neurons (Extended Data Fig. 9**c**). Among all neuron types, relatively more TRAPed tdT+ neurons were found in the mPFC than the BLA (Extended Data Fig. 9**e**).

**Extended Data Figure 9.**
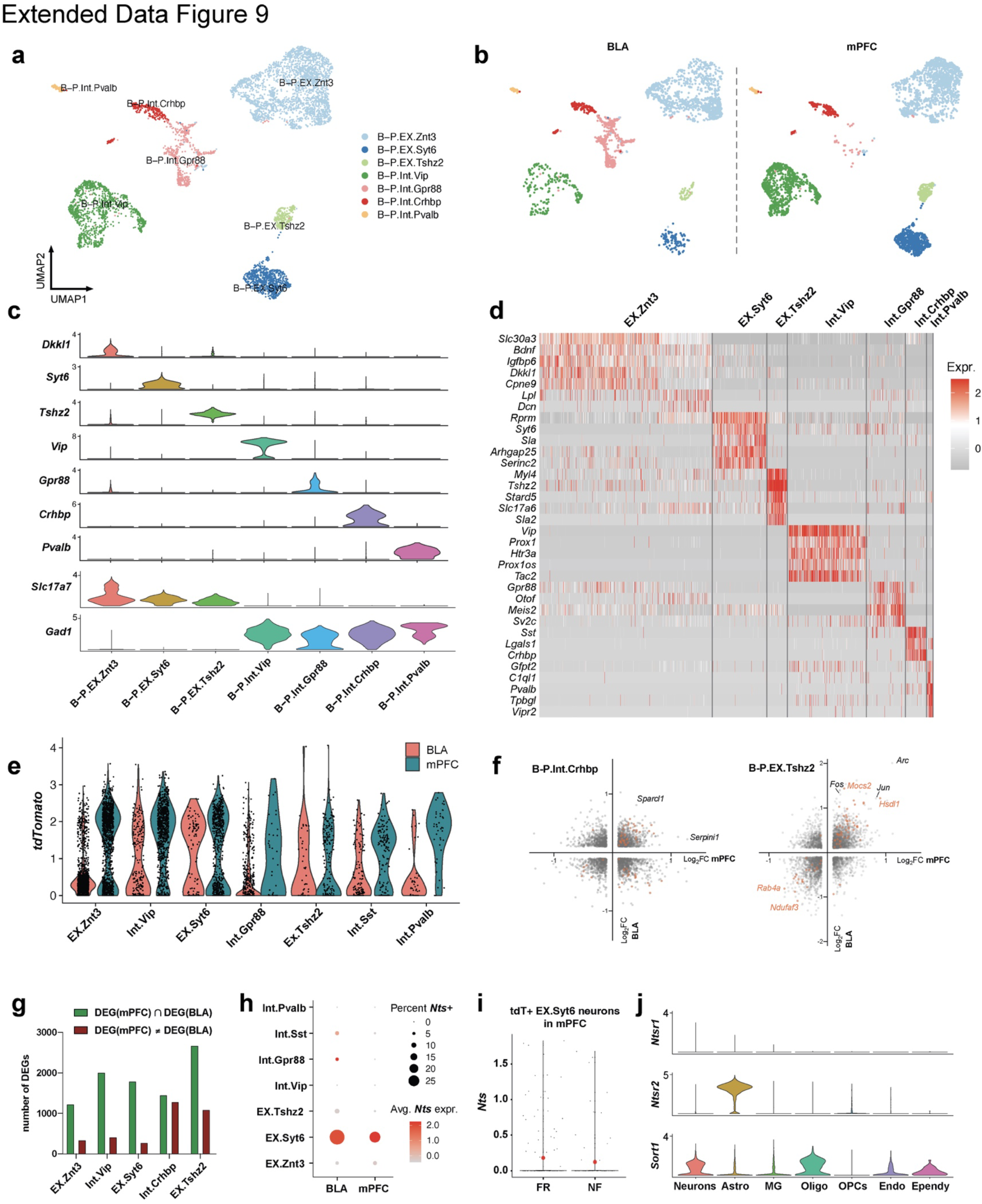
**a)** Integrated clustering of BLA and mPFC neurons. **b)** Integrated clustering of BLA and mPFC neurons separated by regions. **c)** Distinct markers and *Slc17a7* and *Gad1* expression for each cluster of integrated BLA and mPFC clusters. **d)** Heatmap of top marker genes of integrated BLA and mPFC clusters. **e)** tdTomato expression of integrated BLA and mPFC clusters. **f)** DEGs (FR over NF, TRAPed cells) from BLA and mPFC among B-P.Int.Crhbp and B- P.EX.Tshz2 neurons. **g)** Quantification of DEG numbers in each neuron clusters. **h)** *Nts* expression in each neuron clusters in BLA and mPFC **i)** *Nts* expression in tdT+ B-P.EX.Syt6 neurons from mPFC. **j)** Expression of all three known neurotensin receptors in different cell types of BLA.

Next, we examined the FR-induced transcriptional changes within the TRAPed neurons of each type of neuron. Integrated differential expression analysis identified 1,673 genes that were significantly changed in both the BLA and mPFC (Fig. 5**c**, Extended Data Fig. 9**f**). Surprisingly, 1,587 (or 94.9%) of the DEGs were co-regulated in the same direction (Fig. 5**d**). This suggests that memory consolidation drives a highly conserved transcriptional program in engram neurons across multiple brain regions. Consistent with the above analyses, DEGs associated with vesicle exocytosis and synapse formation were upregulated. Finally, within the engram cells of the three most abundant neuron types (EX.Znt3, Int.Vip, and EX.Syt6) we found 32 genes whose expression was consistently modulated by long-term fear memory in both the BLA and mPFC (Fig. 5**e**). Among the top upregulated genes, Polo-like kinase 2 (*Plk2*) is a transcriptional target of *Npas4*^75, 76^ that modulates synapse formation and contextual fear memory,^75^ while Tripartite motif-containing protein 32 (*Trim32*, a E3 ubiquitin ligase^77^), Ubiquitin Like 3 (*Ubl3*), and Ubiquitin C (*Ubc*) are involved in protein ubiquitination that is involved in synaptic plasticity^78, 79^ and fear memory formation in the hippocampus^80^ and amygdala^81, 82^. *Mal2* is an integral membrane constituent of synaptic vesicles associated with vGlut1-positive nerve terminals.^83^ These data suggest that engram neurons in the PFC and BLA share overlapping transcriptional machineries for memory consolidation.

## Neuron-to-astrocyte interaction in remote memory consolidation

In addition to these conserved mechanisms, we also found neurotensin (encoded by *Nts*), a neuropeptide which modulates associative memory in paraventricular thalamus to BLA circuit^84^, was expressed in TRAPed Syt6-positive excitatory neurons (Extended Data Fig. 9**h**) and induced by fear memory consolidation in BLA but not in PFC engram neurons (Fig. 5**f**, Extended Data Fig. 9**i**). This further validates the notion that neuropeptides, including neurotensin, secretogranin, tachykinin, proenkephalin, ProSAAS, and CART, are involved in memory consolidation in BLA engram cells. In addition, neurotensin receptor 2 (*Ntsr2*) is dominantly expressed by astrocytes in the BLA (Fig. 5**g**), while neurotensin receptor 1 (*Ntsr1*) is virtually nondetectable in the BLA and Sortilin (a non-specific neurotensin receptor, encoded by *Sort1*) is detected in multiple cell types in the BLA (Extended Data Fig. 9**j**). *Ntsr2* was shown to be essential for contextual fear memory.^85^ These data suggest that engram neurons in the BLA interact with astrocytes and other cells while consolidating memory.

An atlas of astrocytes across brain regions has described the molecular heterogeneity of astrocytes.^86^ To further understand astrocyte remodeling in memory consolidation, we clustered the integrated data from BLA and mPFC astrocytes 2,278 into four subtypes, in which B-P.A1 cells express thyroid hormone (*Slco1c1*)^87^ and amino acid transporters (*Slc7a10*)^88^, B- P.A2 cells express Calmodulin 1 (*Calm1*) and sphingosine-1-phosphate receptor 1 (*S1pr1*)^86^, B-P.A3 cells express Myocilin (*Myoc*)^89^ and Vimentin (*Vim*)^86^, B-P.A4 cells express synaptic nuclear envelope protein 1 (*Syne1*)^57^, SON DNA and RNA binding protein (*Son*)^86^, and *Utp14b*^58^ (Extended Data Fig. 10**a**–**f**). B-P.A1-3 astrocytes were present in the mPFC and BLA, whereas B-P.A4 astrocytes were specific to BLA (Extended Data Fig. 10**a**,**g**). Furthermore, fear conditioning remodeled the distribution of astrocyte subtypes, in which fear recall induced B-P.A4 in BLA and B-P.A1 in mPFC (Extended Data Fig. 10**g**). Of interest, astrocytes from different training conditions showed consistent *Fos* expression in B-P.A3, but varied *Fos* expression in B-P.A1 and B-P.A4 (Extended Data Fig. 10**b**).

**Extended Data Figure 10.**
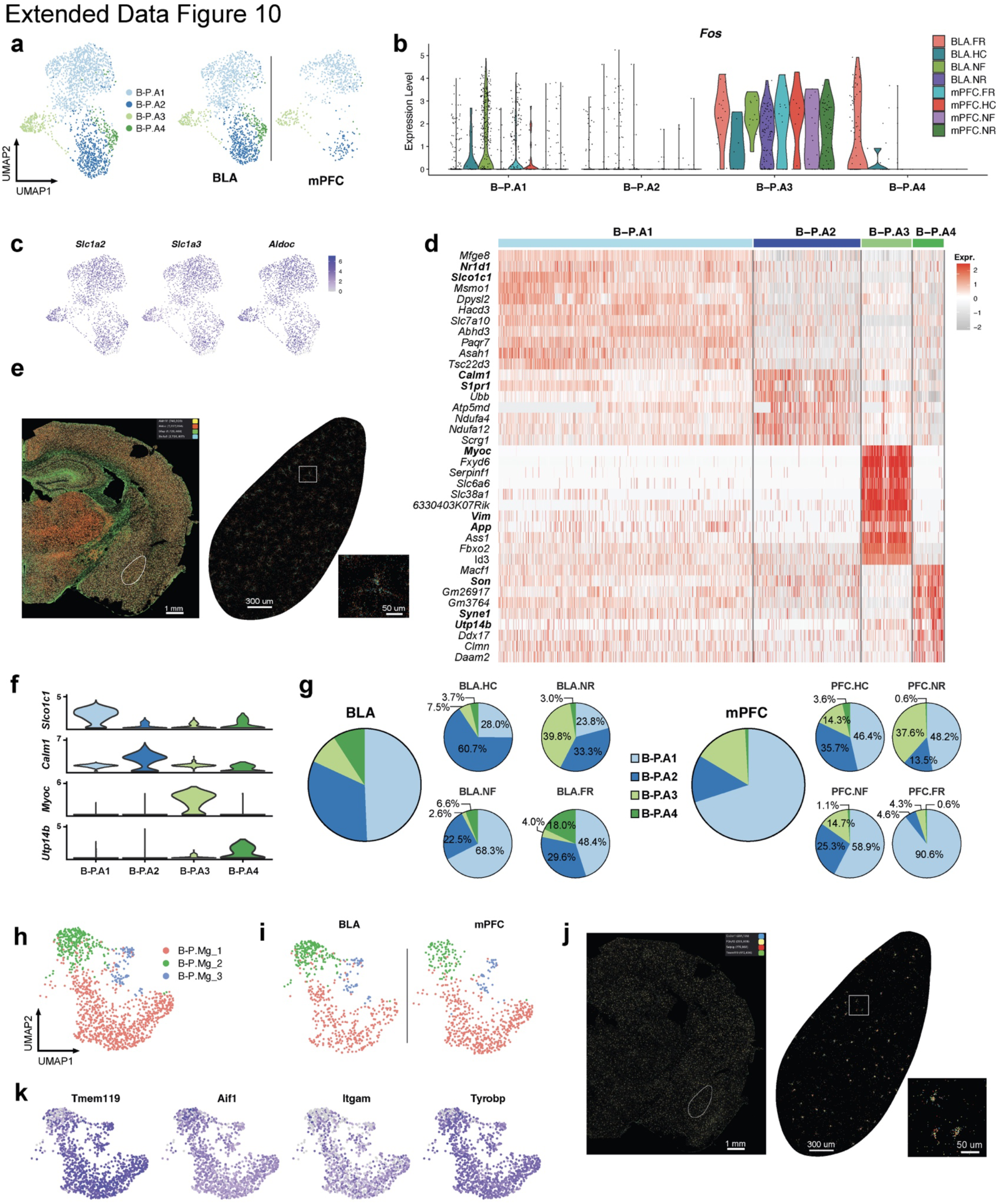
**a)** Integrated clustering of astrocytes from BLA and mPFC single cell RNAseq data. **b)** *Fos* expression separated by astrocyte clusters and condition from BLA and mPFC single cell RNAseq data. **c)** Expression level of astrocyte markers (*Slc1a2*, *Aldoc*, and *Slc1a3*) from BLA and mPFC single cell RNAseq data. **d)** Heatmap of top marker genes of integrated astrocytes cell types from BLA and mPFC single cell RNAseq data. **e)** *Slc1a3*, *Aldh1l1*, *Gfap* and *Aldoc in situ* data from merfish. **f)** Distinct markers expression for each cluster of integrated BLA and mPFC astrocyte clusters, single cell RNAseq data. **g)** Astrocytes compositions in integrated analysis of mPFC and BLA RNAseq, seperated by conditions. **h)** Integrated clustering of microglia from BLA and mPFC single cell RNAseq data. **i)** Integrated clustering of microglia from BLA and mPFC single RNAseq cell data, separated by regions. **j)** *Cx3cr1*, *P2ry12*, *Selplg* and *Tmem119 in situ* data from merfish. **k)** Expression level of pan microglia markers *Tmem119*, *Aif1*, *Itgam*, and *Tyrobp* from integrated BLA and mPFC single cell RNAseq data.

## Summary

Leveraging activity-dependent cell trapping, spatial and single-cell transcriptomics, and chemogenetic experiments, we identified 1) a memory induced activating trajectory of astrocytes; 2) a persistent gene expression program among neurons in mPFC and BLA induced by memory consolidation, independent of salient experience; 3) genes involved in neuronal peptides production, the MAPK pathway, CREB signaling, BDNF signaling, in neuronal synapse assemble, and neuronal projection are upregulated by memory consolidation; 4) fear memory induced Penk and reduced Tac1 in a BLA specific Gpr88+ neurons; 5) a spatially resolved ensemble of engram cells, including *Flt1*+ active astrocytes. The current data provide a solid step towards decoding the comprehensive network of engram cells in long-term memory consolidation.

## Acknowledgements

We thank Drs. E. Jerison and L. Tan for discussion of the experimental design; Dr. N. Neff, V. Tran, A. Seng, and R. Yan for assistance with Vizgen and sequencing; Dr. L. Luo for the gifting and help with TRAP2 line; S.R.Q. is a Chan Zuckerberg Investigator. T.C.S is an HHMI Investigator. The work was supported, in part, by the Swiss National Science Foundation (SNSF 211053 to W.S. and SNSF 211011to H.D.).

## Author Contributions

WS, SRQ, and TCS designed experiments. WS performed animal experiments, brain dissociation, single-cell library preparation, RNAscope, merfish experiment and data analysis. ZL performed brain dissection and virus injection. XJ and MBC contributed part of single cell RNAseq raw data. HD performed flow cytometry. JL performed cell segmentation analysis for merfish data. WS, TCS, and SRQ wrote the manuscript.

## Data and materials availability

All data associated with this study will be available in open repository. Materials are available upon reasonable request.

## Inclusion & ethics statement

We, the authors of this manuscript, recognize the importance of inclusion and ethical considerations in scientific research. Our work is guided by the principles of fairness, transparency, and respect for human dignity.

We affirm our commitment to promoting diversity and inclusivity in science, recognizing that diverse perspectives, backgrounds, and experiences enrich research and enhance scientific discovery. We have made efforts to ensure that our study is conducted in a manner that respects and includes individuals of all races, ethnicities, genders, sexual orientations, abilities, and other aspects of human diversity.

We have obtained all necessary ethical approvals and have followed appropriate guidelines and regulations for the research conducted. We have taken measures to protect the privacy and confidentiality of research participants, including obtaining informed consent and ensuring data security.

We acknowledge the potential for harm in scientific research and have taken steps to minimize any potential harm to research participants or others affected by our work. We have carefully considered the potential implications of our research and have taken responsibility for ensuring that our work is conducted in a manner that upholds ethical and moral standards.

We recognize that scientific research has the potential to impact society in profound ways and we are committed to engaging in responsible research practices that promote the well-being of individuals and society as a whole.

In summary, we affirm our commitment to inclusive and ethical research practices and recognize our responsibility to conduct research that is conducted with integrity, respect, and social responsibility.

## Methods and Materials

### Mice

All animal experiments were conducted following protocols approved by the Administrative Panel on Laboratory Animal Care at Stanford University. The *TRAP2*: *Ai14* mouse line was kindly gifted by the Luo laboratory at Stanford. TRAP2^16^ mice were heterozygous for the *Fos^2A-iCreER^* allele, and homozygous for Ai14 in the C57BL/6 background. Mice were group-housed (maximum five mice per cage) on a 12 h light–dark cycle (07:00 to 19:00, light) with food and water freely available. Male mice 42–49 days of age were used for all the experiments. Mice were handled daily for 3 days before their first behavioral experiment. The animal protocol #20787 was approved by Stanford University APLAC and IACUC. All surgeries were performed under avertin anesthesia and carprofen analgesia, and every effort was made to minimize suffering, pain, and distress.

## Genotyping

The following primers: TCC TGG GCA TTG CCT ACA AC (forward), CTT CAC TCT GAT TCT GGC AAT TTC G (reverse), and ACC CTG CTG CGC ATT G (reporter) were used for genotyping of the *Fos^2A-iCreER^* allele; CTG AGC TCA CCC ACG CT (forward), GGC TGC CTT GCC TTC TCT (reverse), ACT GCT CAC AGG GCC AG (reporter) for wild type allele; CGG CAT GGA CGA GCT GTA (forward), CAG GGC CGG CCT TGT A (reverse), and AAT TGT GTT GCA CTT AAC G (reporter) were used for genotyping of the Rosa- Ai14 allele; TTC CCT CGT GAT CTG CAA CTC (forward), CTT TAA GCC TGC CCA GAA GAC T (reverse), and CCG CCC ATC TTC TAG AAA G (reporter) for Rosa wild type allele.

## Fear conditioning

The fear conditioning training was conducted according to previously described methods^15^. Each mouse was placed individually in the fear conditioning chamber (Coulbourn Instruments), which was positioned at the center of a sound-attenuating cubicle. Prior to each session, the chamber was cleaned with 10% ethanol to provide a background odor, while a ventilation fan generated background noise at around 55 dB. The training began with a 2- minute exploration period, after which the mice received three tone-foot shock pairings separated by 1-minute intervals. Each tone, an 85 dB 2-kHz sound, lasted for 30 seconds, and was followed by a 2-second foot shock of 0.75 mA, with both ending simultaneously. Following each pairing, the mice remained in the chamber for an additional 60 seconds before being returned to their home cages. For context recall, the mice were reintroduced to the original conditioning chamber for 5 minutes, 16 days after the training. 4-OHT injections were administered immediately prior to the recall experiments, within 30 minutes. In the HC and NR groups, 4-OHT was injected at a similar time to the other two groups during the recall. The behavior of the mice was recorded and analyzed using FreezeFrame software (version 4; Coulbourn Instruments), with motionless bouts lasting over 1 second being considered as freezing. Data were analyzed with tracking software Viewer III (Biobserve).

## Brain tissue dissociation and flow cytometry

Basal lateral amygdala was microdissected using a live vibratome sections (300 μm thick). Tissue pieces were enzymatically dissociated using a papain-based digestion system (LK003150, Worthington). Briefly, tissue chunks were incubated with papain (containing L-cysteine), DNase I, and kynurenic acid for 1 hour at 37 °C and 5% CO2. After incubation, tissues were triturated with 300 um glass pipette tips, then 200 um glass pipette tips, and 100 um glass pipette tips. Cell suspensions were then centrifuged at 350g for 10 minutes at room temperature, resuspended in 1 ml EBSS with 10% v/v ovomucoid inhibitor, 4.5% v/v Dnase I, and 0.1% v/v kynurenic acid, and centrifuged again. The supernatant was removed, and 1 ml ACSF was added to the cells. ACSF contained 1 mM KCl, 7 mM MgCl2, 0.5 mM CaCl2, 1.3 mM NaH2PO4, 110 mM choline chloride, 24 mM NaHCO3, 1.3 mM Na ascorbate, 20 mM glucose, and 0.6 mM sodium pyruvate. Cells were then passed through a 70-μm cell strainer to remove debris. Hoechst stain (1:2,000; H3570, Life Technologies) was added and incubated in the dark at 4 °C for 10 minutes. Samples were centrifuged (350g for 8 minutes at 4 °C) and resuspended in 0.5 ml of ACSF and kept on ice for flow cytometry. Live cells were sorted using the BD Vulcan into 384-well plates (Bio-Rad) directly into lysis buffer, oligodT, and layered with mineral oil. After sorting, the plates were immediately snap frozen until reverse transcription.

## Sequencing

Smartseq3 protocol was used for whole-cell lysis, first-strand synthesis, and cDNA synthesis, as previously described with modifications. Following cDNA amplification (23 cycles), the concentration of cDNA was determined via Pico Green quantitation assay (384-well format) and normalized to 0.4 ng/µl using the TPP Labtech Mosquito HTS and Mantis (Formulatrix) robotic platforms. In-house Tn5 were used for cDNA tagmentation. Library were amplified using Kapa HiFi. The libraries were then sequenced on a Novaseq (illumina), using 2 × 100- bp paired-end reads and 2 × 12-bp index reads, with an average of 2 million reads per cell.

## Bioinformatics and data analysis for single cell RNAseq

Sequences from Nextseq or Novaseq were demultiplexed using bcl2fastq, and reads were aligned to the mm10 genome augmented with ERCC (External RNA Controls Consortium) sequences, using STARsolo 2.7.9a. We applied standard algorithms for cell filtration, feature selection and dimensionality reduction. Briefly, genes that appeared in fewer than five cells, samples with fewer than 2000 genes and samples with less than 50,000 reads were excluded from the analysis. Out of these cells, those with more than 10% of reads as ERCC or more than 20% mitochondrial were also excluded from analysis. Counts were log-normalized and then scaled where appropriate. Canonical correlation analysis (CCA) function from the Seurat^90^ package was used to align raw data from multiple experiments. The top 20 canonical components were used. After alignment, relevant features were selected by filtering expressed genes to a set of 2,000 with the highest positive and negative pairwise correlations. Genes were then projected into principal component space using the robust principal component analysis. DEG analysis was done by applying the Mann Whitney Wilcoxon test on various cell populations.

To find memory induced genes in each type of neurons, series of strict criteria were applied. First, we removed the background activation by excluding the DEGs resulted from FR vs. NF among tdT negative neurons. This guarantees their specificity that DEGs are activity-dependent, rather than a general increase in all cells caused by experience. Second, DEGs must be differentially expressed when FR TRAPed cells are compared to NR and HC controls, ensuring that the DEGs were unique to neuronal ensembles associated with memory recall, and not a result of baseline activity (HC) or activity remaining from the initial fear learning (NR). Lastly, each DEG had to meet the criteria of being expressed in a quater of cells and exhibiting at least a 1.75-fold change. By adhering to these standards, a total of 107 DEGs were recognized as “remote-memory-associated DEGs” across six distinct neuron types, BLA.Int.Pvalb was not included in the analysis due to insufficient numbers of cells. EnrichR was used for GO/KEGG/REACTOME pathway analysis and classification of enriched genes (log2FC > 0.5 and *P* <0.05) in each subpopulation.

Single cell RNAseq data from mPFC cells were mapped to mm10 genome with full length tdTomato construct (including Woodchuck Hepatitis Virus Posttranscriptional sequence included in Ai14 line^91^), which improved the sensitivity in calling tdT+ cells. Data from BLA and mPFC cells were integrated using CCA. TRAPed neurons from the each integrated population were analyzed, except B-P.Int.Pvalb and B-P.Int.Gpr88 neurons, due to limited cell number. DEGs with *P* < 0.05 (Mann Whitney Wilcoxon test) were considered as significant DEGs (highlighted in orange Fig. 3**c**, Extended Data Fig. 6**f**).

After unbiased clustering astrocytes, RNA velocity^52^ and Monocle3^53^ were applied to infer astrocytic trajectory. DEGs between FR and NF conditions were estimated using Mann Whitney Wilcoxon test on each astrocyte population.

## RNAscope

The RNAscope multiplex fluorescent reagent kit v2 (323100, ACD) and RNAscope 4-Plex probes were used to conduct the RNAscope experiment according to the manufacturer’s guidelines. The probes employed were either obtained from available stocks or specially created by ACD.

## Gene selection for MERFISH measurements

We used a combination of single-cell RNA sequencing data and literature to select genes for MERFISH. Our selection criteria involved identifying cell-type-marker genes for a particular cell population using a one-vs-all approach. To do this, we performed a Mann Whitney Wilcoxon test for each gene between the cells within the cell population and all other cells not in that population, and corrected the resulting p-values for multiple hypothesis testing to obtain FDR-adjusted p-values. A gene was considered a cell-type marker for a specific cell population if it met the following criteria: 1) expressed in at least 30% of cells within the specified population; 2) FDR-adjusted p-value <0.001; 3) gene expression in the specified population was at least 4-fold higher than the average expression in all cells not in that population; 4) expressed in a fraction of cells within the specified population that was at least two times higher than any other population of cells. We then sorted the marker genes for each population by fold change in expression relative to cells outside the population, and saved the top five marker genes for each population to use for marker selection. In addition to these markers, known genes related to microglias, astrocytes, and OPCs from the literature and included. Finally, DEGs from remote memory associated genes were added to the panel with a total number of 158 genes.

## Tissue processing for MERFISH and RNAscope

Brain tissue samples were processed using a fixed-frozen protocol for both MERFISH and RNAscope. Briefly, animals were sacrificed using CO2 and perfused with cold 4% PFA. Brain tissue were dissected and followed by incubation at 4°C in 4% PFA over-night, 15% sucrose for 12 hours, and 30% sucrose until sink. Brain tissue were frozen in OCT by dry ice and stored in -80 C until sectioning. Sectioning was performed on a cryostat at −18°C. Slices were removed and discarded until BLA region was reached.

Slices with 10 μm in thickness were captured onto Superfrost slides for RNAscope and MERSCOPE slides for MERFISH. The same anatomical region was identified for imaging post hoc after sample preparation, before the start of RNAscope or MERFISH imaging.

## Sample preparation and MERFISH imaging

Slides with tissue sanctions were processed according to MERSCOPE protocol (Vizgen). Briefly, slides with tissue sections were washed three times in PBS, and then stored in 70% EtOH at 4°C for at 18 h to permeabilize the tissue. Tissue slices from the same mouse were cut at the same time and distributed onto four coverslips. After permeabilization, the samples were removed from 70% EtOH and washed with Sample Prep Wash Buffer (PN 20300001), then incubate with Formamide Wash Buffer (PN 20300002) at 37°C for 30 minutes. Gene Panel Mix (RNA probes) were incubated for 48 hours at 37°C. After hybridization, the samples were washed in Formamide Wash Buffer for 30 min at 47°C for a total of two times to remove excess encoding probes and polyA-anchor probes. Tissue samples were then cleared to remove lipids and proteins that contribute fluorescence background. Briefly, the samples were embedded in a thin 4% polyacrylamide gel and were then treated with Clearing Premix (PN 20300003) for 36 h at 37°C. After digestion, the coverslips were washed in Sample Prep Wash Buffer for two times and stain with DAPI/PolyT mix for 15 min. Slides were washed with Formamide Wash Buffer followed by Sample Prep Wash Buffer before imaging. Finally, slides were loaded to MERSCOPE Flow Chamber and imaged at both 20x and 63x magnification.

## MERFISH data processing

MERFISH imaging data were processed with MERlin^92^ pipeline with cell segmentation using CellPose^93^, a deep learning-based cell segmentation algorithm, which based on Hoeschst staining. Decoding molecules were then assigned to the segmented nuclei to produce a cell-by-gene matrix that list the number of molecules decoded for each gene in each cell. The MERFISH expression matrix for each sample was concatenated with the normalized, log-transformed with Scanpy^94^ and integrated using Harmony^95^ and Leiden^96^ clustering was applied to separate the cells into distinct clusters. TRAPed neurons were assigned based on tdTomato expression. DEGs from a comparison of FR-TRAPed and NF-TRAPed conditions were estimated using Mann Whitney Wilcoxon test. Periengram cells were computed as follows: for each engram cell (tdT+), its periengram cells were counted within a radius of 30μm.

## CalEx injection and behavioral experiments

Adeno-associated viruses (AAVs) conveying CalEx^69^ or tdTomato were generated by Addgene based on the vector pZac2.1-GfaABC1D-mCherry-hPMCA2w/b (AAV5, Addgene 111568) or pZac2.1 gfaABC1D-tdTomato (AAV5, Addgene 44332). Stereotaxic procedure of viral microinjection has been described previously. In brief, mice with fear training (within 12 hours) were anesthetized and placed onto a stereotaxic frame (model 1900, KOPF). Mice were injected with Carprofen (5 mg/kg) subcutaneously before surgery. AAVs carrying hPMCA2w/b (CalEx) or control (tdTomato) viruses were loaded via a glass pipette connected with a 10 μl Hamilton syringe (Hamilton, 80308, US) on a syringe injection pump (WPI, SP101I, US) Beveled glass pipettes (1B100–4; World Precision Instruments) filled with viruses were placed into the BLA (1.3 mm posterior to the bregma, 3.4 mm lateral and 3.4 mm medial to the midline, and 4.7 mm from the pial surface). Either 0.3 μl of AAV5 GfaABC1D mCherry-hPMCA2w/b (7e12 vg/ml) or 0.3 μl AAV5 GfaABC1D tdTomato (7e12 vg/ml) were injected at 100 nl/min. Glass pipettes were withdrawn after 10 min and scalps were cleaned and sutured with sterile surgical sutures. Mice were allowed to recover in clean cages for 7 days. Behavioral experiments (recall) were performed at three weeks after surgeries.

## Open field

Mice were placed in the center of 40 × 40 cm white box and allowed to freely explore for 15 min. Videos were recorded and analyzed by BIOBSERVE III software. The 20 × 20 cm region in the center was defined as central zone. The total distance travelled and the activity exploring the center area were analyzed to evaluate the subject’s locomotor ability and anxiety levels.

